# Longitudinal single-cell transcriptional dynamics throughout neurodegeneration in SCA1

**DOI:** 10.1101/2021.10.22.465444

**Authors:** Leon Tejwani, Neal G Ravindra, Billy Nguyen, Kimberly Luttik, Changwoo Lee, John Gionco, Kristen Kim, Jennifer Yoon, Fatema Haidery, Hannah Ro, Luhan Ni, Harry T Orr, Laura PW Ranum, Vikram G Shakkottai, Phyllis L Faust, David van Dijk, Janghoo Lim

## Abstract

Neurodegeneration is a protracted process involving progressive changes in myriad cell types that ultimately result in neuronal death. Changes in vulnerable neuronal populations are highly influenced by concomitant changes in surrounding cells, complicating experimental approaches to interrogate the simultaneous events that underlie neurodegeneration. To dissect how individual cell types within a heterogeneous tissue contribute to the pathogenesis and progression of a neurodegenerative disorder, we performed longitudinal single-nucleus RNA sequencing of the mouse and human spinocerebellar ataxia type 1 (SCA1) cerebellum, establishing continuous dynamic trajectories of each population. Furthermore, we defined the precise transcriptional changes that precede loss of Purkinje cells and identified early oligodendroglial impairments that can profoundly impact cerebellar function. Finally, we applied a deep learning method to accurately predict disease state and identify drivers of disease. Together, this work uncovers new roles for diverse cerebellar cell types in SCA1 and provides a generalizable analysis framework for studying neurodegeneration.

## INTRODUCTION

SCA1 is a progressive neurodegenerative disorder in the family of polyglutamine (polyQ) diseases and is caused by trinucleotide repeat expansion mutations in the CAG tract of *ATXN1*, encoding the ataxin-1 protein (Orr et al., 1993). Although *ATXN1* is ubiquitously expressed (Servadio et al., 1995), the Purkinje cells display the most marked degeneration in the SCA1 cerebellum (Koeppen, 2005) and thus, have been the primary focus of most studies. Evidence from animal models suggests that profound cerebellar circuit dysfunction precedes overt Purkinje cell loss (Watase et al., 2002), which can be attributed to Purkinje cell-intrinsic changes or from perturbations in the many cell types that directly or indirectly modulate Purkinje cell activity. Recent studies have suggested the involvement of additional cerebellar cell types in disease (Cvetanovic et al., 2015; Edamakanti et al., 2018; Kim et al., 2018); however, the effects of SCA1 mutations on other populations are not well-understood and are further obscured by their low relative abundance compared to cerebellar granule cells.

In SCA1 and other neurodegenerative diseases, efforts to interrogate the roles of individual cell types in disease have primarily relied on the development of conditional animal models by restricting mutant gene expression to a candidate cell type, which, in the case of late-onset neurodegenerative diseases, can be extremely laborious and time-consuming. In recent years, droplet-based single-cell sequencing has emerged as a scalable tool for profiling many cells of heterogenous tissues (Habib et al., 2017; Macosko et al., 2015), and several groups have applied this to human post-mortem samples from patients with neurodegenerative disorders (Al-Dalahmah et al., 2020; Grubman et al., 2019; Jakel et al., 2019; Lee et al., 2020; Mathys et al., 2019; Schirmer et al., 2019; Zhou et al., 2020). While these studies have revealed some insights into how certain cell types are unexpectedly perturbed, they reflect the end-stage transcriptional changes in surviving populations of cells, falling short of accurately describing the progressive course of disease, which occurs over decades in humans. Additionally, the mechanisms through which various cell types contribute to disease are constantly changing. Therefore, approaches geared towards elucidating the molecular dynamics throughout a disease process at a single-cell resolution are desirable. Here, we integrate cutting-edge genomics and analytical approaches to examine the complex interplay between diverse cell populations in the SCA1 cerebellum over time.

## RESULTS

### snRNA-seq of the SCA1 cerebellum

To comprehensively profile the transcriptional dynamics of the SCA1 cerebellum throughout the disease at a single-cell level, we performed snRNA-seq of control (CTRL) and SCA1 primary human post-mortem cerebellar cortex tissue (Table S1), and SCA1 *Atxn1^154Q/+^* KI and wild-type (WT) littermate control mice at five timepoints (5, 12, 18, 24, and 30 weeks of age). We selected these timepoints in order to capture all major processes throughout disease, from the early onset of behavioral deficits (5-6 weeks) through the progression to eventual cell loss and premature lethality, which typically begins around 32 weeks of age (Jafar-Nejad et al., 2011; Watase et al., 2002). Following removal of cells that did not meet quality control criteria (see Methods), our snRNA-seq yielded datasets of 41,150 and 318,312 high-quality single-nucleus transcriptomes from human and mouse, respectively (Figure 1A; Table S2). To appropriately merge data across individual samples within a species, we selected batch balanced k nearest neighbors (BBKNN) based on its scalability for large datasets and its ability to integrate data while preserving nuanced biological variation (Luecken et al., 2020; Polański et al., 2019). Finally, to determine the identities of all cells within the human and mouse datasets, we used graph-based unsupervised clustering to generate pre-clusters, imputed missing data using MAGIC (van Dijk et al., 2018) within each genotype (Figure S1), and assigned cell type labels based on expression of established marker genes, merging pre-clusters with similar identities when appropriate (Figures 1B-1E). The resource presented here describes the single-nuclei transcriptomes of a total of 9 human and 13 mouse clusters that represent the major cell types of the cerebellum including granule cells (GC), unipolar brush cells (UBC), Purkinje cells (PC), three classes of GABAergic interneurons (IN1, IN2, and IN3), astrocytes (AS), Bergmann glia (BG), oligodendrocyte progenitor cells (OPC), oligodendrocytes (OL), microglia (MG), pericytes (PER), and endothelial cells (END). Relative proportions of cells and sequencing quality were comparable across genotypes in both the human (Figure 1F; Figures S2A-S2E) and mouse (Figure 1G; Figures S2F-S2I) datasets.

**Figure 1.**
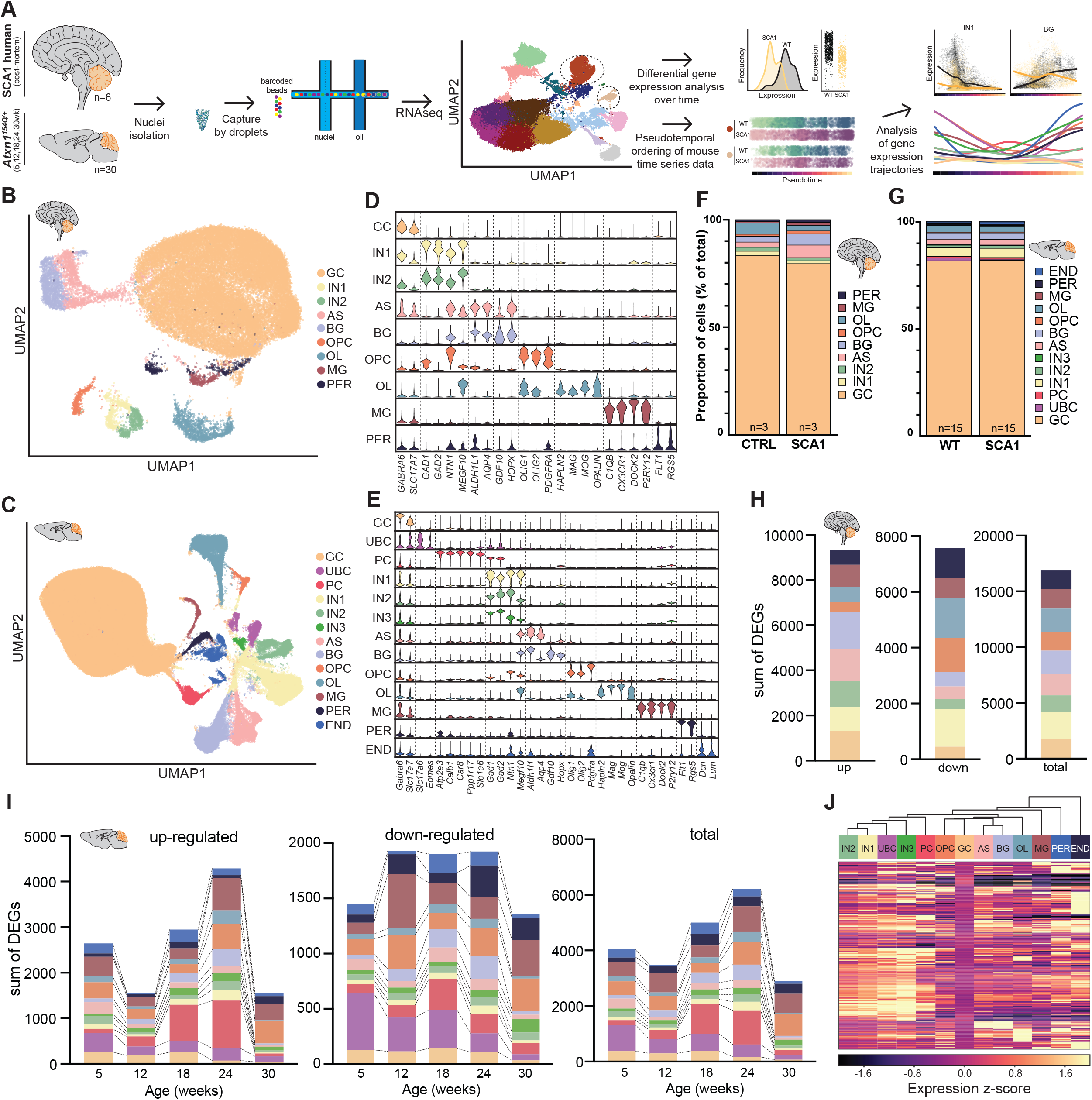
Single-cell transcriptional profiling of the SCA1 human and mouse cerebellum. **(A)** Schematic pipeline of SCA1 snRNA-seq and analysis of human post-mortem (n=6 human) and mouse (n=30 mice) cerebellum. **(B and C)** UMAP embeddings of 41,150 human (B) and 318,312 mouse (C) single-nucleus transcriptomes, which produced the following cell-type clusters: granule cell (GC), unipolar brush cell (UBC), Purkinje cell (PC), three inhibitory interneuron populations (IN1, IN2, IN3), astrocyte (AS), Bergmann glia (BG), oligodendrocyte progenitor cell (OPC), oligodendrocyte (OL), microglia (MG), pericyte (PER), and endothelial cell (END). Nuclei from 6 human post-mortem (CTRL, n=3; SCA1, n=3) and 30 mouse (WT, n=3; SCA1, n=3 per timepoint; timepoints= 5, 12, 18, 24, 30 weeks) tissues were sequenced. **(D and E)** Normalized violin plots showing expression of cell type-specific marker genes for each human (D) and mouse (E) cluster used for cell type annotation. **(F and G)** Relative proportions of each cell type within each genotype for the human (F) and mouse (G) datasets. **(H and I)** Number of up-regulated, down-regulated, and total differentially expressed genes (DEGs) that in the human (H) and mouse (I) snRNA-seq datasets. Genes with imputed |EMD|≥0.1 and P_corrected_<0.01 were considered significant. **(J)** Heatmap displaying mean expression profiles of 237 ataxin-1 physical interactors (rows) in WT mouse cerebellar cell types (columns). *See also* Figures S1 and S2*, and Tables S1-S5.*

### Differential expression of the SCA1 cerebellum

To identify cell type-specific gene expression changes between WT/CTRL and SCA1, we first performed differential gene expression analyses between samples for all cell types and timepoints. Due to the distributional nature of heterogenous gene expression across single nuclei within a population, we computed the Earth Mover’s Distance (EMD) between genotypes for each gene in each cell type, as this approach retained information regarding both the magnitude of change, as well as the proportion of cells undergoing alterations (Nabavi et al., 2016). We identified differentially expressed genes (DEGs) in individual cell types at each timepoint, with different cell types having different numbers of DEGs across the discrete timepoints (Figures 1H and 1I; Tables S3-S5). Overall, the 24-week timepoint had the highest number of DEGs averaged across all cell types in the mouse dataset (Figure 1I; Table S3). Between the different cell types, the PC, OPC, and MG clusters displayed the highest number of DEGs averaged over time (Table S3), with each cluster displaying the highest number of total DEGs at their 24-week timepoint (Figure 1I). Intriguingly, UBCs, an excitatory glutamatergic interneuron population enriched in cerebellar lobules IX and X, the region that undergoes the earliest atrophy in SCA1 patients (Jacobi et al., 2013; Martins Junior et al., 2018), displayed the greatest dysregulation at the 5-week timepoint with regards to the number of DEGs. These data suggest that UBCs, a cell type not previously described in SCA1, may play an unexpected role in early pathogenesis and contribute to regional vulnerability within the cerebellum.

### Expression networks of ataxin-1 interactors

Ataxin-1, the protein mutated in SCA1, is ubiquitously expressed and serves key functions as a transcriptional co-regulator and splicing co-factor through its interactions with various proteins (Zoghbi and Orr, 2009). In SCA1, the majority of gain-of-function pathogenic changes are thought to be mediated through altered incorporation of polyQ-expanded ataxin-1 into its native protein complexes (Lam et al., 2006; Lim et al., 2008; Tejwani and Lim, 2020). Although several key protein interactors of ataxin-1 have been demonstrated to be central to PC pathology, it is unclear if these same molecular mechanisms are responsible for dysfunction of other cerebellar cell types. Thus, we aimed to determine the extent to which various protein interactors could potentially contribute to gene expression changes across the heterogenous cell types of the cerebellum. We hypothesized that for a particular protein interactor to contribute to SCA1 pathogenesis in a given cell type, its simultaneous co-expression with *ATXN1* in the same cell is obligatory. Therefore, we examined the expression of transcripts encoding ataxin-1 interactors on a cluster-by-cluster basis, taking advantage of a comprehensive list of over 200 ataxin-1 physical interactors (Lim et al., 2006). Hierarchical clustering of cell types based on ataxin-1 interactor expression revealed several groups with the most similar expression profiles, including a group of interneuron subtypes (UBC, IN1, IN2, IN3), and an astroglial group (AS, BG), with distinct interactor expression profiles in each of the other cell types (Figure 1J), suggesting that the primary molecular mechanisms governing pathogenesis in different cell types are likely distinct and dependent on co-expression patterns of these interactors with ataxin-1. Because of their high number of DEGs over time (Figure 1I; Tables S3-S5), we focused on analyzing the PC and OPC/OL clusters in greater detail.

### Early synaptic impairments in PCs

In SCA1, while PCs are the primary cell type of the cerebellum to eventually degenerate (Koeppen, 2005), they undergo substantial gene expression changes prior to cell death (Lin et al., 2000). Bulk RNA-seq of the SCA1 mouse cerebellum has revealed several tissue-level changes that have been suggested to be the result of PC perturbations (Driessen et al., 2018; Friedrich et al., 2018; Ingram et al., 2016); however, due to their relative rarity in the heterogenous cerebellum, it remains unclear to what degree this interpretation is confounded by changes in other cell types. Therefore, we sought to conclusively and precisely define the PC-specific dynamics with single-cell accuracy throughout multiple stages of disease for the first time. We first visualized the mouse PC cluster using several dimensionality reduction approaches (Figures 2A and 2B). While standard clustering methods revealed sub-populations within most other cell types, including UBC, IN1, IN2, AS, BG, OPC, OL, MG, and END (Figure S3), they failed to reveal any distinct sub-clusters of PCs (Figure 2A). Additionally, visualization of cell transitions in the PC cluster using PHATE, a dimensionality-reduction method that preserves local and global structure to better display transitions in data (Moon et al., 2019), did not reveal any clear structures in the data that progressed with biological time (Figure 2B). Therefore, we concluded that more nuanced genotype- and time-dependent alterations, rather than global transcriptional changes that would alter positioning in high-dimensional space, were likely to underlie PC dysfunction in SCA1.

**Figure 2.**
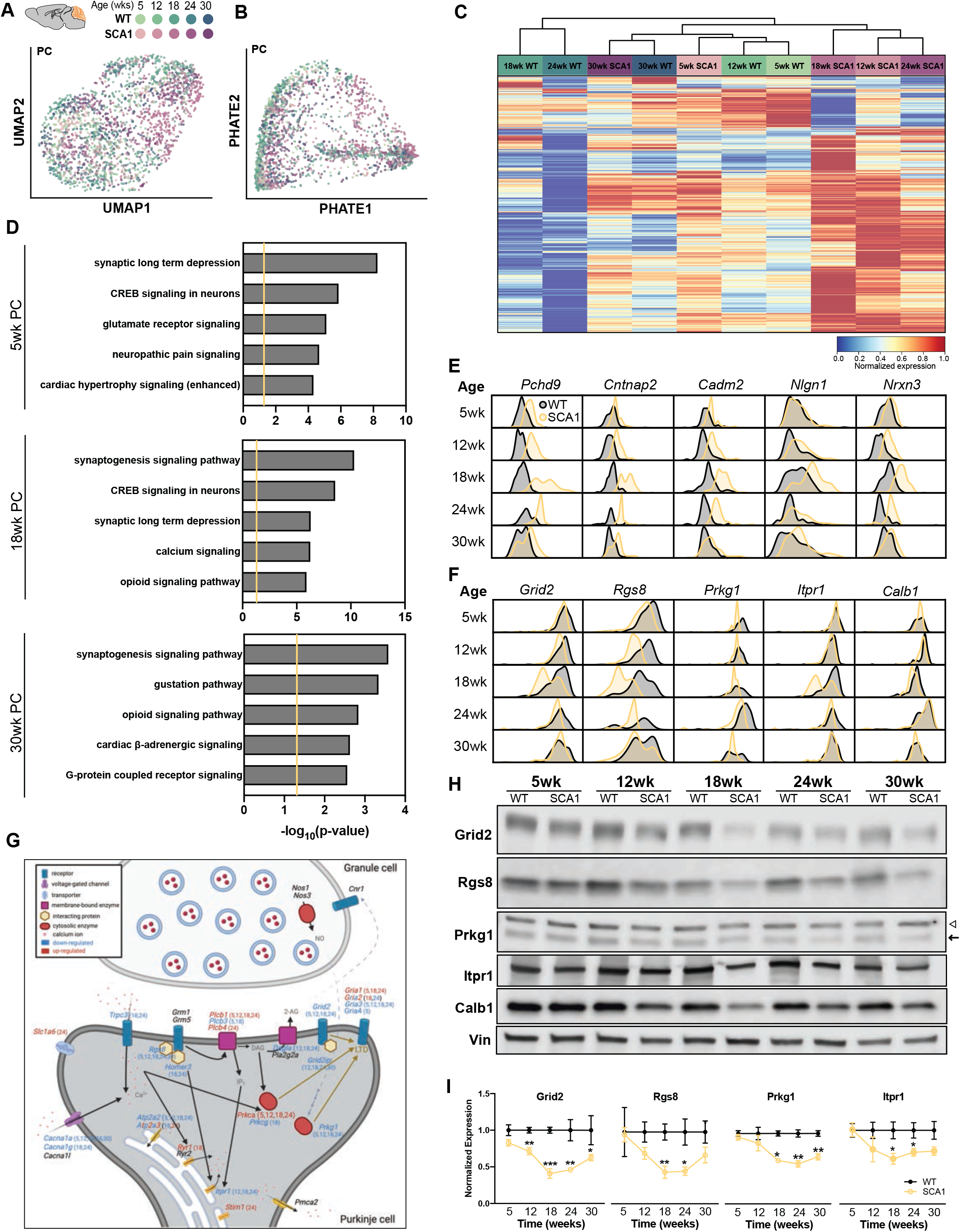
SCA1 Purkinje cell dynamics over time. **(A and B)** UMAP (A) and PHATE (B) embeddings of mouse PC cluster (WT, n=1,152; SCA1, n=1,026 nuclei), colored by genotype and timepoint. **(C)** Heatmap of normalized expression for DEGs (1,578 genes; imputed |EMD|≥0.1 and P_corrected_<0.01 at one or more timepoints) in mouse PCs over time shows the greatest differences in overall gene expression between WT and SCA1 genotypes at 18 and 24 weeks of age. **(D)** Pathway analyses of total DEGs between mouse PCs at 5, 18, and 30 weeks of age. The vertical yellow bar represents a significance cut-off of -log(P-value)>1.3 and the top 5 ingenuity canonical pathways are displayed. **(E and F)** Smoothed histograms displaying time-dependent alterations in distribution of gene expression for selected genes that are up-regulated (E) or down-regulated (F) in SCA1 PCs. **(G)** Summary of gene expression changes at the GC to PC synapse from the mouse snRNA-seq dataset. Information for each DEG regarding direction of differential expression (color) and age (number in parentheses) are provided. **(H and I)** Western blots validating time-dependent changes of several PC-enriched genes/proteins in the 5, 12, 18, 24, and 30 week SCA1 mouse cerebellum (WT, n=4; SCA1, n=4 mice per timepoint). The target band for Prkg1 is denoted with a closed arrow and a non-specific band with an open arrowhead. For data in (I), target expression was normalized to a loading control (vinculin) and then to the average of WT mice at each timepoint. Data presented in (I) are mean±s.e.m. Student’s t-tests were performed to compare genotypes at each timepoint. \**P*<0.05, \*\**P*<0.01, \*\*\**P*<0.001. *See also* Figures S3 and S4*, and Table S6*.

To characterize the progressive transcriptional dysregulation in SCA1 PCs, we next examined differential gene expression in the mouse at 5, 12, 18, 24, and 30 weeks of age. Although gene expression differences were observed in PCs at all timepoints (Figure 1I; Tables S3-S5), hierarchical clustering of WT and SCA1 PCs revealed the largest overall transcriptional divergence between genotypes at 18 and 24 weeks of age (Figure 2C). Surprisingly, of all timepoints sampled, the late-stage 30-week WT and SCA1 PCs clustered most similarly to each other (Figure 2C). This relative normalization of PC gene expression changes at late stages of disease was not only reflected in the number of DEGs, but also in the degree to which DEGs were dysregulated (Figures 1I and 2C; Table S3). Although PC dendritic atrophy is observed at early stages of disease, PC death only occurs at end stages of disease in SCA1 KI mice (Watase et al., 2002); therefore, these data suggest that the eventual PC loss in SCA1 may not depend on the emergence of novel cell-autonomous transcriptional changes in PCs that occur during aging, but rather, involve progressive transcriptional dysregulation in other local populations of cells that communicate with PCs and regulate their function and survival.

We next examined the pathway enrichment for the sets of total, up-regulated, and down-regulated DEGs at each timepoint. Pathway analyses of DEGs in PCs at early timepoints revealed the highest enrichment in genes related to synaptic long-term depression (LTD), CREB signaling in neurons, and glutamate receptor signaling in 5-week PCs (Figure 2D; Table S6). Starting at 12 weeks of age, the most enriched pathway was the synaptogenesis signaling pathway (Figure 2D; Figure S4A; Table S6). Among these DEGs, many cell adhesion and axon guidance signaling molecules involved in synapse formation and maintenance were up-regulated (*Cadm2*, *Cdh12*, *Cntn4*, *Cntn5*, *Cntnap2*, *Dscam*, *Epha5*, *Epha6*, *Epha7*, *Lrrc4c*, *Lsamp*, *Nlgn1*, *Nrxn1*, *Ntrk3*, *Nrxn3*, *Pcdh9*, *Robo2*, *Unc5c*, and *Unc5d*, among others) (Figure 2E; Figure S4A; Tables S5 and S6), likely representing compensatory increases to restore proper synaptic transmission. This early synaptic dysregulation persisted through all timepoints examined. In addition to early synaptic alterations, a persistent down-regulation of genes involved in calcium handling was observed starting from 5 weeks of age (Gene Ontology terms calcium signaling and calcium transport 1; *Atp2a2*, *Atp2b2*, *Cacna1a*, *Cacna2d2*, and *Gria3*) (Figure S4B; Tables S5 and S6), which progressed in magnitude of down-regulation, as well as in number of genes by later timepoints (*Cacna1g*, *Cacnb2*, *Itpr1*, *Trpc3*, among others) (Tables S5 and S6). This early disruption of calcium homeostasis in PCs is consistent with recent studies of SCA1 B05 animals (Chopra et al., 2020), as well as SCA7 transgenic mice (Stoyas et al., 2020). These data suggest that the major PC transcriptional changes are present starting at early stages and are persistently dysregulated throughout the course of disease.

Because impairments in type 1 glutamatergic signaling have been previously suggested to be involved in multiple cerebellar disorders (Kano and Watanabe, 2017), including SCA1 (Friedrich et al., 2018; Power et al., 2016; Serra et al., 2004; Shuvaev et al., 2017), we aimed to define the precise molecular changes in this signaling pathway in PCs. Among all down-regulated protein-coding genes in PCs, the most reduced gene (by EMD) at most timepoints was *Grid2* (Figure S4C; Table S5), encoding the glutamate receptor subunit delta-2 (Gluδ2). Gluδ2 functions as a post-synaptic regulator of mGluR1 specifically at the parallel fiber (PF) to PC synapse (Kato et al., 2012) and is necessary for cerebellar LTD, motor learning, and proper establishment of PF and climbing fiber (CF) territories on PCs (Kashiwabuchi et al., 1995; Kohda et al., 2013). In humans, *GRID2* deletion mutations cause a recessive ataxia (Hills et al., 2013; Utine et al., 2013) and its reduction contributes to cerebellar dysfunction in essential tremor (Pan et al., 2020). Furthermore, loss-of-function *hotfoot* mutations of *Grid2* in rodents (for which over 20 alleles have been described) result in severe ataxia and cerebellar atrophy (Lalouette et al., 1998). Thus, the early and progressive reduction of *Grid2* in SCA1 KI mice is likely to contribute to the onset of motor deficits, as well as the pathological abnormalities in spatial distribution of CF inputs onto PCs observed in SCA1 (Barnes et al., 2011; Duvick et al., 2010; Ebner et al., 2013).

Beyond *Grid2* down-regulation, extensive early changes in other post-synaptic molecules involved in LTD at the PF to PC synapse were also observed at 5 weeks of age, including dysregulation of AMPA receptor subunit-encoding *Gria1*, *Gria3*, and *Gria4*, mGluR1-dependent signal transduction molecules *Plcb1*, *Plcb3*, and *Prkca*, and nitric oxide/cGMP-dependent protein kinase *Prkg1* (Table S5). As disease progressed, additional genes central to cerebellar LTD, PF to PC signaling, and calcium handling were also dysregulated (*Atp2a3*, *Dagla*, *Homer3*, *Grid2ip*, *Itpr1*, *Prkcg*, *Rgs8*, *Ryr1*, *Stim1*, *Trpc3*) (Figures 2F and 2G). Many of these genes were the most differentially expressed at 18 weeks (Figure 2F; Table S5), which we also validated for several PC-enriched genes at the protein level using bulk mouse cerebellar tissue (Figures 2H and 2I). Although PF-evoked synaptic responses in SCA1 PCs are grossly normal (Barnes et al., 2011), the downstream signaling pathways at this synapse described above are critical for the complex functions of the cerebellum, including various forms of plasticity and motor learning; therefore, dysregulation is likely to compromise proper cerebellar functionality. Collectively, these data highlight the PF to PC synapse as a site of early, persistent, and progressive transcriptional dysfunction that coincides with the onset and progression of behavioral impairments in SCA1.

### Myelination is impaired in SCA1

Our snRNA-seq analysis of SCA1 mouse and human post-mortem tissue revealed a large number of DEGs in glial clusters (Figures 1H and 1I; Tables S3-S5), with a high number of DEGs identified in the oligodendroglial lineage (OPC and OL), a glial population suggested to be involved in several other neurodegenerative disorders (Kang et al., 2013; Mot et al., 2018), including at least two other polyQ diseases (Costa et al., 2020; Huang et al., 2015; Ramani et al., 2017). In the central nervous system, OLs serve essential functions in promoting efficient action potential conduction through myelination and in providing metabolic support to axons (Lee et al., 2012; Simons and Nave, 2015). The proper maintenance and plasticity of myelin through adulthood requires continual remyelination, a process in which adult OPCs terminally differentiate into myelin-forming OLs in response to appropriate stimuli (Fields, 2015; Young et al., 2013). Specifically within the cerebellum, myelination of PC axons is necessary for precise synaptic transmission onto deep cerebellar nuclei neurons (Barron et al., 2018), which integrate information from the cerebellar cortex with excitatory inputs from extracerebellar regions, and subsequently project to their target output regions throughout the central nervous system. Furthermore, adult remyelination is essential for motor learning (McKenzie et al., 2014), a major function of the cerebellum that is impaired at early timepoints in SCA1 KI mice (Watase et al., 2002).

Because many significantly down-regulated genes of the human SCA1 OL cluster encode proteins restricted to terminally-differentiated OLs (Figures 3A and 3B), and because there was a large dysregulation of genes in mouse and human OPCs (Table S3), we hypothesized that the formation of mature, myelinating OLs from OPCs may be impaired. To examine the dynamics of the OPC to OL conversion, we estimated the overall RNA velocity of the human oligodendroglial lineage (OPC and OL clusters) and subsequently visualized cell trajectories using PHATE (Figure 3C) (Bergen et al., 2020; La Manno et al., 2018; Moon et al., 2019). Through comparing the relative abundance of spliced and unspliced transcripts, RNA velocity can infer the kinetics of gene expression, providing meaningful insights into the future transcriptional states of differentiating cells (La Manno et al., 2018). This analysis revealed two transcriptionally distinct differentiation courses of *PDGFRA*^+^ OPCs into *MBP*^+^*MOG*^+^ mature OLs in CTRL samples, while SCA1 OPCs only differentiated into OLs along a single trajectory (Figure 3C; Figure S5A), demonstrating the loss of a subset of transitioning cells. Furthermore, differentiated human SCA1 OLs expressed lower levels of mature OL transcripts, including several myelin protein-encoding genes (Figure 3D; Figure S5B). Differential expression of multiple transcription factors involved in OL formation and maturation was observed in the OPC (*MYT1*, *OLIG1*, *OLIG2*, *SOX10*) and OL (*NKX2-2*, *OLIG1*, *OLIG2*, *NKX6-2*, *ZNF24*, *ZNF488*, *ZNF536*, *MYRF*) clusters (Figure S5C), suggesting that transcriptional dysregulation at multiple stages during differentiation may underlie a partial OPC block, as well as a failure to initiate and maintain appropriate expression of pro-myelinating genes.

**Figure 3.**
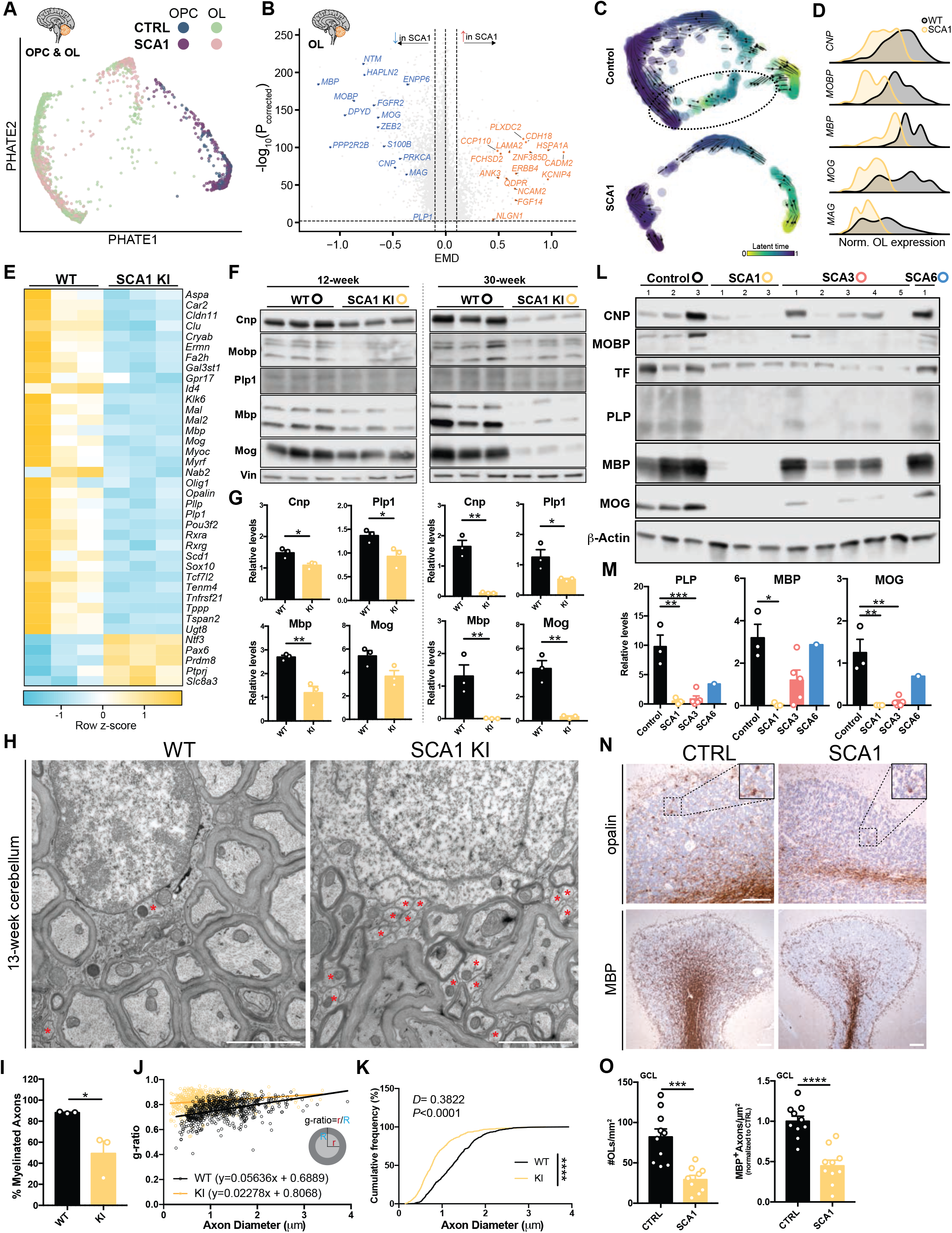
Oligodendrocyte impairments and myelin deficits in SCAs. **(A)** PHATE embedding of human oligodendroglia lineage (OPC+OL clusters; CTRL, n=1,055; SCA1, n=959 nuclei). **(B)** Volcano plot displaying DEGs of the human OL cluster. Genes with imputed |EMD|≥0.1 and P_corrected_<0.01 were considered significant. **(C)** RNA velocity (scVelo) PHATE embeddings of human oligodendroglia lineage to examine OPC to OL differentiation trajectories. The dotted line denotes a population of differentiating OPCs/OLs absent in SCA1. **(D)** Smoothed histograms displaying distribution of expression of major myelin protein-encoding transcripts in the human OL cluster, showing significant decreases in SCA1. **(E)** Heatmap of significantly differentially expressed OPC/OL genes from bulk RNA sequencing of the 12-week WT and SCA1 KI mouse cerebellum (WT, n=3; SCA1, n=3 mice). **(F and G)** Western blots confirming myelin protein reduction in the 12- and 30-week SCA1 KI mouse cerebellum (WT, n=3; SCA1, n=3 mice per timepoint). **(H-K)** Transmission electron microscopy of the 13-week WT and SCA1 KI mouse cerebellum (H) revealed a reduction in percentage of myelinated PC axons (I), increased g-ratio (J), and reduction of axon diameter (K) in SCA1 KI mice. Scale bar=2μm. Red asterisks indicate nonmyelinated axons. Only myelinated axons were quantified in (K). **(L and M)** Validation of myelin protein reduction in the SCA1, SCA3, and SCA6 human post-mortem cerebellar cortex relative to controls (CTRL) (CTRL, n=3; SCA1, n=3; SCA3, n=5; SCA6, n=1 patient). **(N and O)** Immunohistochemistry (N) and quantification (O) of OL number (i.e. opalin-positive cells) and myelination (MBP-positive staining) in the granule cell layer (GCL) of the human post-mortem cerebellar cortex (CTRL, n=10; SCA1, n=10 patients). Opalin, scale bar=100μm. MBP, scale bar=100μm. For bar plots, data are mean±s.e.m. The following statistical tests were used: Student’s t-tests to compare across genotypes for (G, I, and O); one-way ANOVA with multiple comparisons to compare across genotypes for (M); Kolmogorov-Smirnov test to compare frequency distributions in (K). \**P*<0.05, \*\**P*<0.01, \*\*\**P*<0.001, \*\*\*\**P*<0.0001. *See also Figures S5-S7*.

To interrogate the course of oligodendroglial impairments during SCA1 progression, we examined oligodendroglia throughout time in the SCA1 KI animal. Although OPCs and OLs comprise a small fraction of the total cells of the cerebellum (Figures 1F and 1G; Figures S2A, S2B, S2F, and S2G), the consequences of their dysfunction were observed at the tissue-level as early as 12 weeks of age, as demonstrated by bulk RNA-sequencing (Driessen et al., 2018) with qRT-PCR validation (Figure 3E; Figures S6A and S6B), and by biochemical analysis of myelin proteins, which revealed a progressive loss of myelin proteins over time (Figures 3F and 3G). Ultrastructurally, SCA1 KI mice displayed an age-dependent reduction in myelination of PCs (Figures 3H-3J; Figures S6C-S6E) and PC axon diameter (Figure 3K; Figure S6F). Importantly, these myelin-related changes were not secondary to PC dysfunction, as SCA1 PC-specific B05 transgenic mice (*Pcp2-ATXN1[82Q]/+*) did not display the same deficits (Figures S6G-S6J), suggesting the possibility of cell-autonomous impairments in OL function. As expected from the snRNA-seq analyses, a similar reduction of myelin proteins was observed in SCA1 and SCA3 human post-mortem cerebellar tissue, but not SCA6 (Figures 3L and 3M), consistent with neuroimaging studies of patients (Lukas et al., 2006; Mandelli et al., 2007). Human tissue-level changes in myelination were further confirmed by qRT-PCR (Figure S7A) and by immunohistochemistry in the cerebellar cortex of a larger cohort of SCA1 patients (Figures 3N and 3O; Figures S7B-S7I). Collectively, these data demonstrate that myelin deficiencies are a common feature of several SCAs and may exacerbate cerebellar dysfunction through enhancing PC structural and physiological deficits.

### Continuous dynamics throughout disease

Although differential gene expression analysis of time-series data provides a granular view of transcriptional alterations at specific timepoints throughout disease, chronic neurodegenerative disorders involve long-term, progressive changes that are not necessarily able to be categorized into a small number of discrete stages. Furthermore, individual cells sampled at a specific timepoint exist on a continuum of transcriptional cell identities that may resemble those from other timepoints. In order to broadly characterize each cerebellar cell type’s variable contribution to SCA1, we aimed to generate cell trajectories that capture continuous gene expression changes over time. Recently developed trajectory inference methods have been applied to reconstruct complex lineages of varying topologies through pseudotemporal ordering of cells from single-cell datasets that undergo profound transcriptional changes over time, such as during differentiation (Kanton et al., 2019) or cellular reprogramming (Schiebinger et al., 2019). To the best of our knowledge, similar application to chronic biological datasets to assess continuous gene expression dynamics of cells that retain their cellular identities but undergo subtle, but progressive changes over aging, such as during neurodegeneration, has not yet been reported.

To computationally infer cell type-specific gene expression dynamics from our SCA1 KI mouse time-series data, we first ordered single cells using MELD (Burkhardt et al., 2020), which was able to accurately reconstruct overlapping trajectories within each cell type. Overall, these continuous trajectories matched with progression through biological time (Figure 4A), also visualized by PHATE (Figure S8). By doing this, we were able to examine longitudinal gene expression patterns, simultaneously comparing the heterogeneous dynamics of differential expression between genotypes over the course of disease for each cell type. We specifically focused on the top dynamical genes, as these were most likely to be involved in progressive events over aging. To calculate dynamical genes, we used a machine learning model to regress the pseudotime per cell given each cell’s transcriptome, which achieved an average R^2^ of 0.92 across cell types and genotype. Using this approach, we ranked features by information gain, selecting the top 100 WT and 100 SCA1 predictive genes within each cell type (Table S7). By comparing the mean expression difference between WT and SCA1 for each dynamical gene across pseudotime (Figure 4B) and averaging across all top dynamical genes within a cell type, we generated a roadmap of the SCA1 cerebellum that describes the degree of overall perturbation for each cell type throughout all points during disease (Figure 4C). In addition to confirming the results of our differential gene expression analyses regarding early dysfunction of UBCs and OPCs (Figure 1I; Table S3) and the peak dysregulation of PCs at mid-late disease stages (Figure 2), this analysis also provided an overview of the transcriptional dysregulation in all other cell types over time. As an example, targeted examination of several dynamical genes in AS and BG (a specialized type of unipolar AS) revealed varying longitudinal expression profiles in these related cell types (Figure 4D), highlighting the utility of our single-cell approach in precisely understanding diverse gene expression patterns in individual cell types.

**Figure 4.**
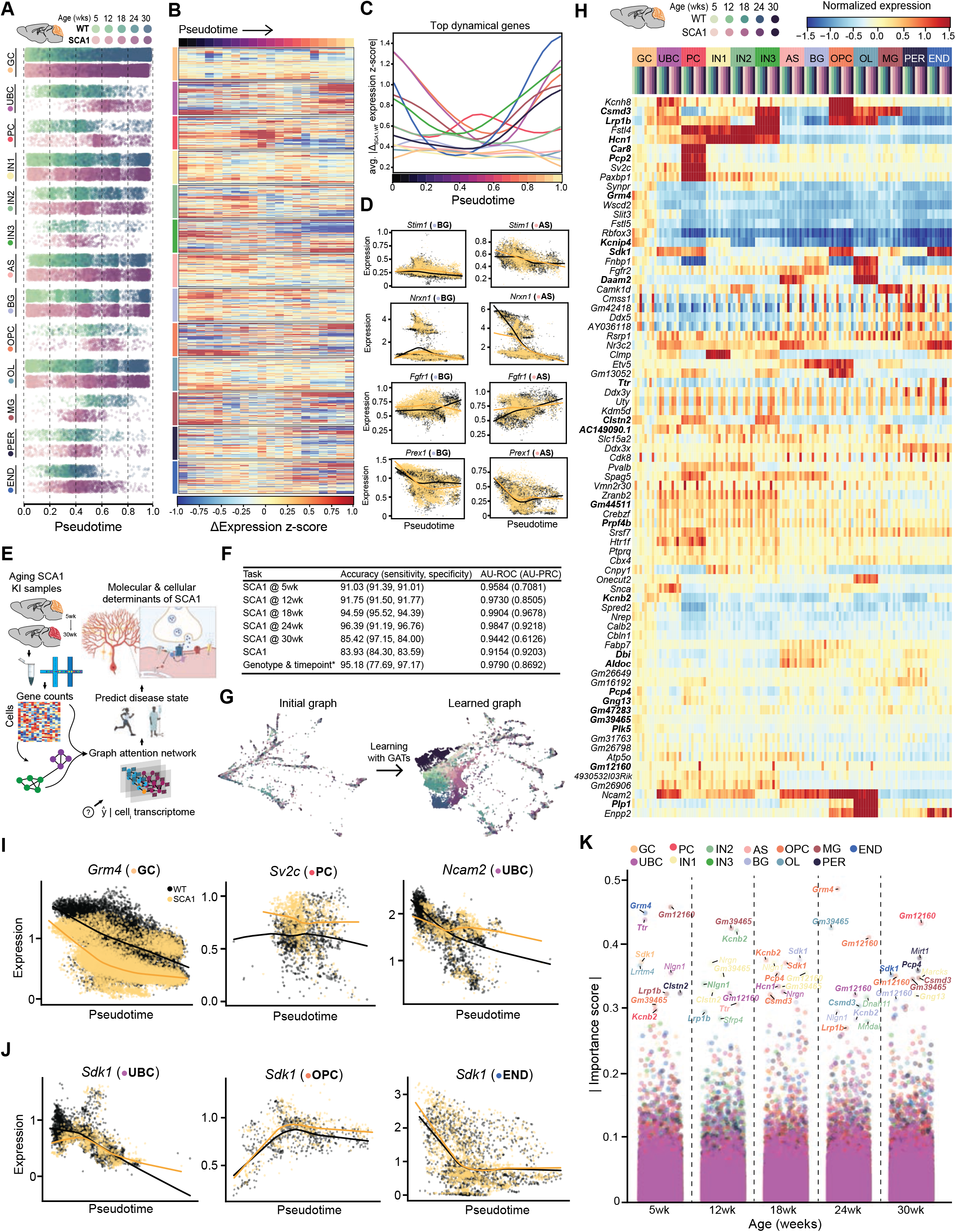
Cell trajectories and deep learning to predict disease state and progression. **(A)** Pseudotemporal ordering of mouse time series data using manifold enhancement of latent dimensions (MELD) to generate continuous progressions for each cell type cluster. **(B)** Heatmap displaying Δ_SCA1-WT_ expression z-score for top 100 WT and 100 SCA1 dynamical genes within each cell type over pseudotime. **(C)** Average |Δexpression| between genotypes of the top 100 WT and 100 SCA1 dynamical genes within each cell type over pseudotime to highlight timing of each cell type’s dysregulation. Expression was represented as z-score (removal of the mean and dividing by the standard deviation across all cells) to standardize the displayed units. **(D)** Expression dynamics of select dynamical genes over pseudotime show differential patterns between BG (left) and AS (right) clusters. **(E)** Schematic overview of approach using geometric deep learning on snRNA-seq data to predict disease state at single-cell resolution in order to gain insights into pathophysiology. **(F)** Performance and evaluation of each model on the test set across 6 binary tasks and 1 multi-label classification task (*) (AU-ROC=area under the receiver-operating characteristic curve, AU-PRC=area under the precision-recall curve). For the multi-label classification task, performance metrics were averaged across classes after prevalence-weighting in a one-vs.-rest heuristic evaluation scheme. **(G)** PHATE embeddings of test set cells based on the original cell-similarity graph (left), which served as input to each GAT model, and the graph learned by one of the attention heads in the GAT model discriminating WT from SCA1 cells (right). **(H)** Heatmap of z-score of mean expression of top 20 salient genes per attention head in the 5 binary classification tasks predicting SCA1 at various timepoints. Z-score of expression was averaged for each genotype, timepoint, and cell type, where the mean was calculated from the first moment of each gene’s expression across the test set. **(I and J)** Expression dynamics of select salient genes from (H) over pseudotime shows that the GAT model pays attention to genes that are highly differentially expressed between genotypes (I), as well as genes with more nuanced patterns of differential expression (J). **(K)** Importance scores for each gene across various cellular subsets. Importance was calculated by computing Integrated Gradients, with a SmoothGrad noise tunnel, for the average expression of each cell subset across the test set, relative to a baseline of no expression. The top 10 important genes for each timepoint are labeled. Bolded genes in (H and K) are salient/important for both the GAT and Integrated Gradient approaches. *See also Figure S8, and Tables S7-S9.*

### Deep learning to predict disease drivers

Conventional differential gene expression analyses rely on gene-by-gene comparisons to associate genes with disease but may fail to identify interaction effects between genes, as well as marginal signals that, although weaker, are nonetheless associated with disease (Cordell, 2009; Wang and Blei, 2020). On the other hand, deep artificial neural networks can be used to identify complex non-linear interactions between input features; however, deep networks are challenging to train and difficult to interpret. To better understand the complex dependencies between a cell’s transcriptome, developmental trajectory, and cell type, we trained several models on our longitudinal SCA1 snRNA-seq data and applied a new approach to interpret these black-box models in order to extract deeper insights into the transcriptional patterns of SCA1.

The first challenge of any deep network is to train, i.e., to learn, how to classify cells given their transcriptome and generalize to unseen data with good performance accuracy. To do this, we used Graph Attention Networks (GATs), a geometric deep learning architecture, to train models in an inductive setting across 6 binary and 1 multi-label classification tasks (Figure 4E). Graph neural networks rely on a message-passing scheme, where features are aggregated according to a graphical structure and a new representation of the input graph and features are learned by gradient-based optimization and backpropagation. In predicting whether an individual cell exhibits transcriptional signatures of SCA1 at each timepoint, this approach yielded a minimum area under the receiver-operating characteristic curve (AU-ROC) of 0.94, suggesting that our models learn to detect unique SCA1 signatures at various timepoints (Figure 4F). In addition, this approach allowed us to take a single cell, of any timepoint or cell type, and determine with ∼80% accuracy (AU-ROC, 0.92) whether the cell exhibits a SCA1 transcriptional signature. As such, we can use deep learning to determine SCA1 status at any point during the lifetime of the mouse at the organismal scale with high accuracy, as well as to detect cells that exhibit early signs of SCA1 within cerebellar tissue. Finally, to demonstrate that our architecture can learn the genotypic and developmental variability across the dataset, we trained a graph neural network to discriminate between 10 labels at single-cell resolution. This model was able to discriminate both genotype and age with ∼95% accuracy (macro-averaged, one-vest-rest AU-ROC 0.98), supporting the interpretation of this model to identify the transcripts and cells that distinguish SCA1 from WT (Figure 4F).

GATs build on Graph Convolutional Networks by learning additional coefficients that determine the extent to which the models should attend to each cell’s neighbors when aggregating their features to learn a new, dimensionality-reduced representation of each cell node and predict their labels. The coefficients that each attention head learns can be interpreted as a new representation of the input graph. Since both the node and graph representations are trained by optimizing the objective function, these representations force each cell to attend to others in a way that maximally discriminates cells to yield better label predictions. We aggregated these coefficients for each attention head to construct new weighted graphs, which we used for downstream manifold learning in order to identify phenotypical populations or subsets of cells with distinct transcriptomic signatures. This semi-supervised approach to manifold learning enabled a better separation of cells according to their age and disease status labels (Figure 4G), indicating that the model pays attention to cells according to their genotype and timepoint labels. These new learned embeddings can be clustered to further identify populations of cells for downstream differential analyses and sub-setting based on cell type, which may indicate unique disease states.

After confirming the predictive accuracies of our models (Figure 4F), we next applied local and global algorithms to interpret the features within the dataset to which the models pay the most attention when predicting age and disease status labels, which we refer to as “gene saliency” (Figure 4H; Table S8). We hypothesized that because expression patterns within these genes were crucial for the highly accurate genotype and age label predictions, these genes may represent features that are important to the pathogenesis and/or progression of SCA1. Because GATs learn co-variations in gene expression between all cell types, the nature of an individual genes’ saliency to the models may vary locally, i.e., cells at different distances from the decision boundary may have variable importance weights across the same set of genes. To more specifically interrogate how the top globally salient genes contribute to SCA1 biology, we first examined their expression within individual cell types over time using our longitudinal mouse snRNA-seq dataset (Figure 4H). Among the genes enriched in specific cell types, robust gene expression differences between genotypes were observed for some salient genes within that population (Figure 4I), while for other genes, differential expression in predicted cell types was not apparent (Figure 4J), demonstrating that GATs pay attention to a combination of diverse transcriptional signals.

Examining global gene saliency may omit subtle differences between various types of cells. To overcome this limitation, we applied a recently developed approach in the field of explainable AI (XAI), Integrated Gradients (Sundararajan et al., 2018), to each model in order to attribute the models’ output to the variable importance of input features for cells of different cell types and timepoints. Rather than looking at global model saliency for a single input layer, Integrated Gradients calculates the adjustment that the learned model would make for a given input transcriptome throughout the computation graph. We also added a noise tunnel, which introduces Gaussian noise to the input expression, and averaged across the Integrated Gradients to produce a more robust importance score per input feature. This analysis revealed a second, independent set of important genes, many of which overlapped with top salient genes from the GAT (Figures 4H and 4K; Tables S8 and S9). In addition to validating our earlier deep learning approach, this method revealed the specific cell types and timepoints during which each important gene had the highest importance score (Figure 4K), providing additional granularity and interpretability of this approach. Collectively, these deep learning approaches identified both prominent and nuanced transcriptional signatures that contribute to SCA1 pathogenesis and progression and offer a complementary approach to traditional clustering and differential expression analysis of single-cell data.

## DISCUSSION

Here, we have generated a longitudinal transcriptomic atlas of the mouse and human SCA1 cerebellum that describes the simultaneous transitions of individual cell types within a heterogenous tissue with single-cell precision. These data reveal new insights into the molecular and cellular underpinnings of SCA1 pathogenesis, with broad implications for other cerebellar and neurodegenerative disorders. First, we have comprehensively examined the transcriptional dysregulation in PCs, the major cerebellar cell type lost in SCA1, throughout disease. Surprisingly, we observed a normalization of PC gene expression at late stages in our mouse dataset. Although it is possible that the most severely affected PCs had already degenerated by the 30-week timepoint and that the data reflect the transcriptomes of the less-affected, surviving PCs, this is unlikely, as this timepoint immediately precedes the bulk of cell loss in the SCA1 KI model, suggesting that the progressive, late-stage events leading to eventual PC death likely involve dysfunction of surrounding cell populations that regulate PC survival. Furthermore, we have also detailed the transcriptional changes that affect the PF to PC synapse, a major site of dysfunction in SCA1. In doing so, we have not only validated the findings of earlier studies (Driessen et al., 2018; Friedrich et al., 2018; Ruegsegger et al., 2016; Serra et al., 2004), but have also provided a great degree of additional granularity into the extent and the precise timing of events underlying the impairment of glutamatergic signaling transmission at this synapse. This analysis has uncovered several key molecules for PC function that are dysregulated at early stages, such as Gluδ2, GluA1, GluA3, GluA4, and Prkg1, among others, which, along with changes at CF to PC synapses, may explain the emergence of early behavioral deficits in SCA1.

Next, our data also revealed a profound transcriptional dysregulation in various glial populations, including oligodendroglia. Although OPCs receive bona fide synaptic input from multiple neuron populations to stimulate their terminal differentiation, including PCs of the cerebellum (Karadottir et al., 2008; Zonouzi et al., 2015), our data demonstrate that the OL deficits observed in SCA1 are not strictly secondary to PC dysfunction. Whether the impairments in OL differentiation reported here are a cell autonomous result of mutant ataxin-1 expression in differentiating OPCs or a consequence of dysfunction in additional populations of cells with which they interact, such as CFs from the inferior olivary nucleus (Lin et al., 2005), remains to be determined. Nonetheless, due to their integral role in regulating the metabolic health and survival of axons (Lee et al., 2012), promoting differentiation of OPCs into OLs to increase myelination may provide a novel therapeutic entry point for multiple cerebellar disorders. Beyond PCs and OPCs/OLs, this study uncovers the unique molecular signatures of all major cell types of the cerebellum throughout the course of SCA1 and will serve as an invaluable foundational resource for the future experimental dissection of the complex interplay between these cell types in cerebellar health and disease.

In addition to revealing specific insights into the pathogenesis of SCA1, the analysis pipeline developed here represents an important advance in the integration of single-cell technologies with novel computational tools to better understand dynamic events. By constructing continuous single-cell trajectories of chronic time-series data, we have enabled the examination of progressive gene expression changes within a cell type over the course of disease, overcoming limitations of traditional approaches in which analysis of gene expression differences is restricted to and within a small number of discrete timepoints. We have also applied an xAI approach that not only accurately predicts labels related to cells in the snRNA-seq dataset, but can also be used to interpret the molecular determinants of those labels that underlie the models’ predictive power, going beyond gene-by-gene comparisons at discrete timepoints to include patterns in multi-gene dynamics that concomitantly contribute to disease. Although we have applied this approach to SCA1, a monogenic disorder with a known genetic cause, the application of our xAI approach to similar datasets for idiopathic disorders may reveal the molecular drivers of disease, while also providing diagnostic utility via the identification of early disease signatures. In summary, we believe the pipeline developed here and applied to SCA1 can serve as a prototype for future studies of the dynamics of a variety of large-scale quantitative data in the context of neurodegeneration or any progressive event.

## ACKNOWLEDGMENTS

We thank Dr. Thomas Biederer and all members of the Lim and van Dijk laboratories for useful feedback, critiques, and comments. We are grateful to Matthew Perkins (University of Michigan Brain Bank), Dr. Arnulf H. Koeppen (Veterans Affairs Medical Center, Albany, New York, USA), the New York Brain Bank (Columbia University, New York, New York, USA), and the National Institutes of Health Neuro-BioBank (University of Miami Brain Endowment Bank, Miami, FL, USA) for post-mortem materials and would like to thank all the patients and families that contributed to brain donation. We would also like to thank Dr. Guilin Wang (Yale Center for Genome Analysis) for assistance with snRNA-seq and Dr. Xinran Liu (Yale Center for Cellular and Molecular Imaging) for assistance with electron microscopy. The University of Michigan Brain Bank is supported by the NIH (P01 AG053760). This work was supported by National Institutes of Health grants NS083706 (J.L.), NS088321 (J.L.), MH119803 (J.L.), AG066447 (J.L.), T32 NS041228 (L.T., K.L.), T32 NS007224 (L.T.), the Lo Graduate Fellowship for Excellence in Stem Cell Research (L.T.), the Gruber Science Fellowship (L.T.), the UCSF Medical School Summer Explore Research Fellowship (B.N.), and the Yale College First-Year Fellowship in the Sciences & Engineering (J.Y., H.R.).

## AUTHOR CONTRIBUTIONS

L.T., N.G.R., D.v.D., and J.L. conceived and designed this study. L.T. performed human and mouse snRNA-seq. N.G.R. analyzed the snRNA-seq data, with L.T. providing guidance on biological interpretation. B.N. and L.T. performed validation of Purkinje cell deficits in SCA1 KI mice. L.T., K.L., K.K., F.H., H.R., J.Y, and L.N. performed validation of oligodendrocyte deficits in SCA1 KI mice. L.T. and C.L. performed biochemical validation of myelin proteins SCA human post-mortem tissue. L.T. and K.L performed ultrastructural analysis of SCA1 KI and B05 mouse cerebellum and H.R., J.Y., and L.T. quantified and analyzed the electron microscopy data. J.G. and P.L.F. performed histological studies of oligodendrocyte changes in human post-mortem tissue. H.T.O. provided reagents, critical feedback, and discussion. L.R. and V.S. provided frozen human post-mortem tissue for snRNA-seq and biochemical analyses. N.G.R. and D.v.D. developed the mathematical foundations of the GAT method. L.T., N.G.R., D.v.D., and J.L. wrote the manuscript. All authors edited the manuscript and provided comments.

## DECLARATION OF INTERESTS

The authors declare no competing interests.

## METHODS

• KEY RESOURCES TABLE
• RESOURCE AVAILABILITY

○ Lead Contact
○ Materials Availability
○ Data and Code Availability
• EXPERIMENTAL MODEL AND SUBJECT DETAILS

○ Animal Husbandry
○ Human Tissue Samples and Patient Information
• METHOD DETAILS

○ Isolation of Nuclei from Frozen Tissue
○ 10x Genomics snRNA-seq
○ Protein Extraction and Western Blotting
○ Total RNA Extraction and qPCR
○ Electron Microscopy
○ Human Cerebellum Histology and Microscopy
• QUANTIFICATION AND STATISTICAL ANALYSIS

○ Integration and Clustering of snRNA-seq Data
○ Differential Expression and Pathway Analyses
○ Expression of Ataxin-1 Interactors
○ RNA Velocity
○ Image Analysis
○ Trajectory Analysis
○ Deep Learning of snRNA-seq Data
○ Statistical Analysis

## RESOURCE AVAILABILITY

### Lead Contact

Further information and requests should be addressed to and will be fulfilled by the Lead Contact, Janghoo Lim.

### Materials Availability

This study did not generate new unique reagents.

### Data and Code Availability

All raw fastq data files from human and mouse snRNA-seq were deposited into GEO under accession #xxxxxx. The data and additional details of protocols used for this study are available from the Lead Contact upon reasonable request. Scripts used to analyze the snRNA-seq data and the software for the Graph Attention Networks and Integrated Gradients with noise tunnels are also available at the following link for academic use: https://github.com/xxxxxxxxxx/xxxxxx-xxxx.

## EXPERIMENTAL MODEL AND SUBJECT DETAILS

### Animal Husbandry

All animal procedures were performed in accordance with the National Institutes of Health Guide for Care and Use of Experimental animals and approved by the Yale University Institutional Animal Care and Use Committee. SCA1 knock-in (SCA1 KI; *Atxn1^154Q/+^*) and PC-specific transgenic (SCA1 B05; *Pcp2-ATXN1[82Q]/+*) mice, along with their littermate controls, were maintained on a pure C57BL/6J background. Mice were maintained on a 12 hours light and 12 hours dark cycle with standard mouse chow and water *ad libitum*. For animal PCR genotyping, tail snips for DNA extraction were taken for between P18-P20 for all animals. Both male and female mice were used for biochemical and sequencing analyses (see Table S2). Mouse tissue for biochemical analysis was collected as previously described (Driessen et al., 2018).

### Human Tissue Samples and Patient Information

Frozen human post-mortem tissue used for snRNA-seq, qRT-PCR, and western blotting was provided by the Michigan Brain Bank through Dr. Vikram Shakkottai and The Center for NeuroGenetics (Florida) through Dr. Laura Ranum. For histological analyses, ten human SCA1 cerebellar samples were obtained from a hereditary ataxia specimen repository at the Veterans Affairs Medical Center in Albany, New York (courtesy of Dr. A.H. Koeppen). Six age-matched control cerebellar samples were from the New York Brain Bank, acquired from prospectively followed individuals that were clinically free of neurodegenerative disorders on serial neurological examinations through the Alzheimer’s Disease Research Center or the Washington Heights Inwood Columbia Aging Project at Columbia University, and subsequently confirmed with neuropathological examination of the post-mortem brain (Vonsattel et al., 2008). Four control brains were obtained from the National Institutes of Health NeuroBioBank (University of Miami, Miami, FL, USA). Standardized measurements of brain weight (grams) and post-mortem interval (hours between death and placement of brain in a cold room or upon ice) were recorded. Brains from male and female donors were used for all analyses. All individuals signed informed consent forms approved by the respective university or institutional ethics boards.

## METHOD DETAILS

### Isolation of Nuclei from Frozen Tissue

Nuclei were isolated from frozen tissue as previously described (Habib et al., 2017). For mouse snRNA-seq, three animals per genotype (WT and SCA1 KI) per timepoint (5, 12, 18, 24, and 30 weeks) were used, for a total of 30 animals. For human post-mortem samples, comparable regions of cerebellar cortex were harvested from frozen tissue of three samples per diagnosis (CTRL and SCA1), for a total of six human samples. Briefly, frozen mouse or human tissue was gently dounce homogenized in 2mL of ice-cold Nuclei EZ Prep buffer (Sigma, Cat. NUC101-1KT) with large clearance pestle “A” and then a small clearance pestle “B” for 25 strokes each, then incubated on ice for five minutes with an additional 2mL of cold EZ Prep buffer. Nuclei were centrifuged at 500xg for five minutes at 4°C. Pelleted nuclei were then washed in 4mL of cold EZ Prep buffer, incubated on ice at five minutes, and centrifuged again at 500xg for five minutes at 4°C. Pelleted nuclei were then washed in 4mL of Nuclei Suspension Buffer (NSB) containing 1x PBS, 0.01% BSA, and 0.1% RNase inhibitor (Clontech/Takara, Cat. 2313B) and centrifuged at 500xg for five minutes at 4°C. Finally, nuclei were resuspended in 1mL of NSB, filtered through a 40μm cell strainer (Fisher Scientific, Cat. 22-363-547), and counted using a Countess II FL Cell Counter (Applied Biosystems, Cat. A27978). Single-nucleus suspensions were diluted to approximately 1000 nuclei per μl in NSB.

### 10x Genomics snRNA-seq

Libraries were prepared from diluted single-nucleus suspensions using the 10x Genomics Chromium Single Cell 3’ Reagent Kits v.3 (mouse dataset) or v.3.1 (human dataset) through the Yale Center for Genome Analysis (YCGA). Briefly, 10,000 cells per sample were added to RT Master Mix and loaded on the Single Cell A Chip and portioned with a pool of about 750,000 barcoded gel beads to form nanoliter-scale Gel Beads-In-Emulsions (GEMs). Each gel bead has primers containing the following: an Illumina R1 sequence (read 1 sequencing primer), a 16-nucleotide barcode, a 12-nucleotide unique molecular identifier (UMI), and a 30-nucleotide poly-dT primer sequence. Upon dissolution of the Gel Beads in a GEM, the primers were released and mixed with the cell lysate and Master Mix. Incubation of the GEMs then produced barcoded, full-length cDNA from poly-adenylated mRNA. Silane magnetic beads were then used to remove leftover biochemical reagents from the post-GEM reaction mixture.

Full-length, barcoded cDNA was then amplified by PCR to generate sufficient mass for library construction, and enzymatic fragmentation and size selection were used to optimize the cDNA amplicon size prior to library construction. P5, P7, a sample index, and R2 (read 2 primer sequence) were added during library construction via end repair, A-tailing, adaptor ligation, and PCR. The final libraries contained the P5 and P7 primers used in Illumina bridge amplification. Libraries were sequenced using the NovaSeq6000 at the YCGA.

Sample concentrations were normalized to 1.2nM and loaded onto an Illumina NovaSeq S4 flow cell at a concentration that yielded 20k-50k passing filter clusters per cell per sample. Samples were sequencing using paired-end sequencing on an Illumina NovaSeq6000 instrument according to Illumina protocols using 10x sequencing specifications. The 8bp index was read during an additional sequencing read that automatically followed the completion of read 1. Data generated during sequencing runs were simultaneously transferred to the YCGA high-performance computing cluster. A positive control (prepared bacteriophage Phi X library) provided by Illumina was spiked into every lane at a concentration of 0.3% to monitor quality in real time. Signal intensities were converted to individual base calls during a run using the system’s Real Time Analysis (RTA) software. Base calls were transferred form the machine’s dedicated personal computer to the Yale high-performance computing cluster via a 1 Gigabit network mount for downstream analysis.

### Protein Extraction and Western Blotting

For all biochemical validation experiments, tissues from both male and female mice of each genotype were used. Total protein lysates were obtained from frozen mouse or human cerebellar tissue by dounce homogenization in Triple lysis buffer (0.5% NP-40, 0.5% Triton X-100, 0.1% SDS, 20mM Tris-HCl [pH 8.0], 180mM NaCl, 1mM EDTA, Roche cOmplete protease inhibitor cocktail, and PhosStop protease inhibitors), rotated for 15 minutes at 4°C, and centrifuged for 10 minutes at 13,000rpm at 4°C. The supernatant was isolated and total protein concentration was measured with the BCA protein assay kit (ThermoFisher). Equal protein amounts were boiled at 95°C for 10 minutes and analyzed by SDS-PAGE (BioRad). Separated proteins from gels were transferred for one hour at 100V onto nitrocellulose membranes at 4°C. Membranes were then washed with TBST (Tris-buffered saline, 0.1% Tween-20) three times for ten minutes each, blocked by 5% non-fat dry milk in TBST for one hour at room temperature, followed by incubation with primary antibody in 5% non-fat dry milk in TBST at 4°C overnight. On the next day, membranes were washed with TBST three times for ten minutes each and incubated with sheep anti-mouse or donkey anti-rabbit IgG-conjugated with horseradish peroxidase (HRP) (GE Healthcare; 1:4,000) for 90 minutes at room temperature. Membranes were then washed with TBST three times for ten minutes and developed using SuperSignal West Pico Plus Chemiluminescent substrate (Pierce, Cat. 34580) and visualized using a KwikQuant Imager (Kindle Biosciences).

The following primary antibodies were used: rabbit anti-GRID2 (Abcam; ab198499; 1:5,000), rabbit anti-ITPR1 (Invitrogen; PA1-901; 1:1,000), rabbit anti-PRKG1 (MyBioSource; MBS1493952; 1:1,000), rabbit anti-RGS8 (Novus Biologicals; NBP2-20153; 1:1,000), mouse anti-Calbindin-D-28K (Sigma-Aldrich; C9848; 1:1,000), mouse anti-CNPase (Abcam; ab6319; 1:200), rabbit anti-MOBP (Bioss; bs-11184R; 1:300), rabbit anti-Plp1 (Abcam; ab28486; 1:1,000), mouse anti-MBP (Abcam; ab62631; 1:1,000), mouse anti-MOG (Millipore; MAB5680; 1:1,000); mouse anti-Vinculin (Sigma-Aldrich; V9264; 1:10,000), rabbit anti-transferrin (Abcam; ab82411; 1:1,000); and mouse anti β-Actin (Sigma; A5316; 1:10,000).

### Total RNA Extraction and qPCR

RNA was extracted from mouse cerebellum or comparable regions of frozen human post-mortem tissue using the Qiagen RNeasy Mini Kit as recommended by the manufacturer’s instructions. For quantitative RT-PCR (qRT-PCR), cDNA was synthesized using oligo-dT primers and the iScript cDNA synthesis kit (BioRad). qRT-PCRs were performed using Taqman probes with iTaq Universal Probe Supermix on a C1000 Thermal Cycler (BioRad) equipped with BioRad CFX Manager software. The following Taqman probes (Applied Biosystems) were used: *CNP (*Hs00263981_m1)*, MOBP* (Hs00197813_m1), *PLP1* (Hs00166914_m1)*, MBP* (Hs00921945_m1)*, MOG* (Hs01555268_m1), *ACTB* (Hs01060665_g1), *GAPDH* (Hs_02758991_g1), *HPRT1* (Hs02800695_m1), *Mbp* (Mm01266402_m1), *Plp1* (Mm01297210), *Actb* (#4352933E), and *Hprt* (Mm03024075_m1). Expression data were determined by normalizing target expression to housekeeping genes (*ACTB*, *GAPDH*, and *HPRT1* for human; *Actb* and *Hprt* for mouse) using BioRad CFX Manager software and plotted using Prism 8 (GraphPad).

### Electron Microscopy

Mice were anesthetized with isoflurane and transcardially perfused with 1x PBS for one minute, 4% paraformaldehyde (PFA) in PBS for 4 minutes, and then in 1x PBS for one additional minute. Whole brains were removed and fixed in EM fixative solution 1 (2.5% glutaraldehyde and 2% PFA in 0.1M sodium cacodylate pH 7.4) overnight at 4°C. The cerebellum was sectioned coronally in cold PBS using a vibratome (Leica VT1000) at a thickness of 150μm. Sections were further post-fixed in EM fixative solution 2 (1% OsO4 and 0.8% potassium ferricyanide in 0.1M sodium cacodylate pH 7.4) for one hour at room temperature, then washed three times for 5 minutes in EM rinse buffer (0.1M sodium cacodylate pH 7.4). Specimens were then stained *en bloc* with 2% aqueous uranyl acetate for 30 minutes, dehydrated in a graded series of ethanol to 100%, substituted with propylene oxide, and embedded in EMbed 812 resin. Sample blocks were polymerized in an oven at 60°C overnight. Thin 60nm sections were cut with a UC7 ultramicrotome (Leica) and post-stained with 2% uranyl acetate and lead citrate. Sections were examined with a FEI Tecnai transmission electron microscope at 90kV accelerating voltage and digital images were recorded with an Olympus Morada CCD camera and iTEM imaging software at the Yale University CCMI Electron Microscopy Facility.

### Human Cerebellum Histology and Microscopy

Formalin fixed tissue was embedded in paraffin and 7μm-thick cerebellar sections were stained with luxol fast blue-hematoxylin and eosin (LH&E). Purkinje cells were counted as previously described (Choe et al., 2016) and number of Purkinje cell axonal torpedoes in the entire section was normalized by dividing the Purkinje cell layer length to account for variations in the amount of cerebellar cortex in the tissue block (expressed as mm^-1^).

For immunohistochemistry, 7μm-thick paraffin-embedded cerebellar sections were rehydrated and antigen-retrieval was performed with Trilogy solution (Sigma-Aldrich) in a conventional steamer for 40 minutes, followed by a 20-minute cooldown at room temperature. Endogenous peroxidase was inactivated by incubation in 3% H2O2. Sections were blocked for two hours at room temperature in 10% normal goat serum, 1% BSA, 0.1% Triton X-100 in PBS, pH 7.4, and then incubated overnight at 4°C in primary antibody. The next day, after several washes in PBS-0.1% Tween, sections were incubated with goat anti-mouse (1:500) or goat anti-rabbit (1:200) biotinylated secondary antibodies (Vector Labs), followed by peroxidase substrate (Vectastain ABC kit) and developed with 3,3’-diaminobenzidine in Dako DAB+ Chromogen solution (Agilent, Cat. K3468). The following primary antibodies were used: mouse anti-MBP (Biolegend; SMI-99P; 1:5,000) and rabbit anti-opalin (Abcam; ab121425; 1:100). Brightfield microscopic images were acquired blindly on an Olympus Bx43 microscope with a Lumenera Infinity 3 digital camera.

## QUANTIFICATION AND STATISTICAL ANALYSIS

### Integration and Clustering of snRNA-seq Data

Following alignment of sequencing reads to the pre-mRNA-containing mouse or human reference genomes (mm10 or GRCh38, respectively) using the cellranger count function (10x Genomics), we followed the standard scRNA-seq analysis pipeline for pre-processing scRNA-seq count data (Satija et al., 2015). All snRNA-seq data were analyzed with Python (version 3.8.2) and pre-processing was performed using SCANPY v1.6.0 (Wolf et al., 2018). Briefly we removed genes that were expressed in fewer than 3 nuclei and removed nuclei that expressed fewer than 200 genes prior to imputation, as well as nuclei in which greater than 10% of the total counts aligned to mitochondrial genes. The resulting UMI counts were normalized to library size and square-root transformed.

To integrate snRNA-seq samples from different samples within a species, we compared various integration methods (i.e. Seurat v3, COMBAT, LIGER, Mutual nearest neighbors, and Harmony) (Haghverdi et al., 2018; Korsunsky et al., 2019; Polański et al., 2019; Stuart et al., 2019; Welch et al., 2019) and ultimately used an approximate batch-balanced KNN (BBKNN) graph for batch-effect correction, manifold learning, and clustering, which allowed the quick and accurate generation of the two mouse and human snRNA-seq pooled datasets (Luecken et al., 2020). We utilized SCANPY’s fast approximation implementation of BBKNN with 100 principal components to calculate pairwise Euclidean distances but otherwise default parameters (Polański et al., 2019; Wolf et al., 2018). BBKNN assumes that at least some cell types are shared across batches and that differences between batches for the same cell type are lower than differences between cells of different types within a batch. For each cell, the 3 nearest neighboring cells in each condition were identified by Euclidean distance in 100-dimensional Principal Component Analysis (PCA) space. This kNN graph was used as the basis for downstream analysis.

To visualize the snRNA-seq data, we implemented various non-linear dimension reduction methods with the BBKNN batch-corrected connectivity matrix as input for uniform manifold approximation and projection (UMAP) and potential of heat-diffusion for affinity-based trajectory embedding (PHATE) (Becht et al., 2018; Moon et al., 2019). UMAP projections were generated using a minimum distance of 0.5 and PHATE projections were generated with a gamma parameter of 1. For clustering nuclei, we used the Louvain community detection method applied to the BBKNN graph with high resolution (resolution=3, otherwise default parameters using SCANPY’s implementation), which facilitated cell type annotation. We used the BBKNN graph for gene expression imputation using Markov affinity-based graph imputation of cells (MAGIC) (van Dijk et al., 2018). Then, pre-clusters of the same major cerebellar cell type were merged when appropriate and manually annotated based on imputed expression of previously described cell type-specific marker genes to yield 1 of 13 cell types.

### Differential Expression and Pathway Analysis

To determine differentially expressed genes, we used a combination of three metrics: the Wasserstein or Earth Mover’s distance (EMD), an adjusted *P*-value from a two-sided Mann-Whitney U test with a continuity correction and a *P*-value adjustment using the Benjamini-Hochberg correction, and the binary logarithm of fold change between mean counts. The EMD can be defined as the minimal cost to transform one distribution to another, and has been used to assess gene expression that significantly differs between conditions (Orlova et al., 2016; Wang and Nabavi, 2018). We performed binary comparisons of genotype for each timepoint and cell type for imputed expression (batch-effect corrected and conditioned on genotype) and for normalized and transformed raw data (data not shown) separately. Genes with *P*_corrected_< 0.01 and |EMD|≥0.1 were considered significant. Finally, pathway enrichment analyses of significantly differentially expressed genes were performed using Ingenuity Pathway Analysis (QIAGEN Inc., https://www.qiagenbioinformatics.com/products/ingenuitypathway-analysis). Enriched canonical pathways were considered significant above a threshold of [-log(*P*-value)>1.3].

### Expression of Ataxin-1 Interactors

Ataxin-1 protein interactors have been previously identified using a yeast two-hybrid approach (Lim et al., 2006). The comprehensive list of these interactors and several others was obtained from the Biological General Repository for Interaction Datasets (BioGRID) (Stark et al., 2006). To visualize the cell type-specific expression of these ataxin-1 interactors, we first standardized expression for each gene at each timepoint, removing the average expression across cell type and genotype and dividing by the standard deviation (z-score). We then averaged the z-score in each cell type and hierarchically clustered the expression data with a Euclidean distance metric for agglomeration using scipy (version 1.5.2). The resulting averaged, z-scored expression for all interactors in WT cerebellar cells was plotted in a heatmap.

### RNA Velocity

To analyze the splicing dynamics and transcriptomic regulation of oligodendroglia (OPC and OL clusters) in human post-mortem samples, we aligned reads to segregate spliced and unspliced counts (La Manno et al., 2018). From these layered count matrices, we computed the RNA velocity for individual cells using scVelo, a generalization of the original RNA velocity method that allowed for characterization of multiple transcriptional states (Bergen et al., 2020). We calculated the first and second order moments with 50 principal components and assumed 30 neighbors. We then fit a stochastic RNA velocity model with default parameters. We visualized the learned dynamics, stratified by diagnosis, and the relation of spliced and unspliced counts using scVelo and our original embeddings.

### Image Analysis

For electron microscopy studies, 20-25 images were obtained for each animal and quantified blindly using the Fiji package of ImageJ. Following de-identification of images, percentage of myelinated axons in each image was manually quantified using the Cell Counter plugin and averaged across all acquired images within an animal. The longest outer myelin diameter and overlapping inner myelin diameter (axon diameter) were measured and the *g*-ratio was calculated by dividing the inner myelin diameter by the outer myelin diameter. Axons were excluded if microtubule organization appeared non-orthogonal to the sectioned plane being imaged.

For quantitative histology studies of human post-mortem cerebellar tissue, whole slide scanning was performed on a Leica Biosystems Aperio AT2 scanner and analyzed with Aperio ImageScope software (v.12.4.0). Areas of cerebellar cortex were outlined, the number of MBP-labeled profiles were quantified using the positive pixel count v9 algorithm, and cells in the granule cell layer with a distinct rim of cytoplasmic opalin staining were quantified with the counting function. The area of white matter and total cerebellar cortex were outlined on MBP immunostained sections in StereoInvestigator software (MicroBrightfield) using the tracing function.

### Trajectory Analysis

To quantify the temporal ordering of cells, we relied on the extent to which an individual cell is transcriptionally similar to a known timepoint sample, inferring pseudotime based on a graph signal processing approach called Manifold Enhancement of Latent Dimensions (MELD) (Burkhardt et al., 2020). We encoded a raw experimental score for each cell in the dataset such that -1, -0.5, 0, 0.5, and 1 corresponded to cells from 5, 12, 18, 24, or 30wk mice, respectively. Using the kernel from the BBKNN graph (described above), these raw scores were smoothed in the graph domain, yielding a pseudotime per cell. We visualized these results by plotting expression against the pseudotime and fitting a smooth line to these data, using locally estimated scatterplot smoothing or local regression (LOESS) with two-thirds of the data in each of the three residual based re-weightings, as implemented with statsmodels (version 0.11.1). This method can be thought of as a linear trajectory for a particular gene across a particular timepoint, where pseudotime is calculated for each cell type independently.

To determine which genes drive the dynamics of transcriptomic changes in WT and SCA1 mice across development and aging, we used a machine learning approach to regress pseudotime given gene expression. Namely, for each cell type and genotype, we took the cell by genes feature matrix and used it as input to a model to predict each cell’s pseudotime, 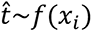 where, *f* represents a machine learning model; in our case, we used a gradient boosting tree model to approximate the inferred time per cell *i*, given its gene expression *x*. To improve predictive performance, evaluated as mean squared error of the predicted cellular time compared to the inferred pseudotime, we imputed data conditioned on genotype, achieving an average (*R*^2^= 0.92 across cell type, genotype, and cross-validation folds. To implement our gradient boosting tree model, we used XGBoost (Python implementation, version 1.2.0) (Chen and Guestrin, 2016).

We used Bayesian optimization with a Tree Parzen estimator to select hyperparameters that maximized performance (Bergstra et al., 2013). Briefly, we used a learning rate of 0.3, a max tree depth of 100, a minimum Hessian needed in a child of 5, 80% subsampling of training instance per boosting iteration, and a 25% sample of features used for each tree, leaf, and node. Otherwise, default hyperparameters were used. Each fold was trained for a maximum of 5,000 epochs with a patience of 500 epochs on the cross-validation fold dataset. We used 10-fold cross-validation to select the best model for feature engineering. Using the best model over cross-validation folds per cell type and genotype, we determined dynamical genes by capitalizing on tree-based model’s implicit feature importance algorithm: tree-based models add splits to a tree when there is a wrongly classified element upstream such that each of the branches from a new split are more accurate. Thus, we used information gain to rank which genes are important to predicting inferred time, sorting genes by the improvement in accuracy that each gene feature provides to the branches that it is on in the final model. The top 100 genes most important, by information gain, to predicting a cell’s time, across the training data, were considered to be the top 100 dynamical genes.

### Deep Learning of snRNA-seq Data

Deep learning is able to model non-linear and higher-order dependencies between features within a dataset. As such, we developed a deep learning approach and applied it to our mouse snRNA-seq dataset to identify transcriptomic determinants of disease that may have escaped detection in differential gene expression analyses. Graph Attention Networks (GAT) have been shown to achieve much better performance on single-cell datasets than other models, which suggested that we could use this graph neural network (GNN) architecture to learn to discriminate transcriptomic signatures of SCA1 from WT at single-cell resolution (Ravindra et al., 2020; Sehanobish et al., 2020). GATs have a self-attention mechanism that builds on graph convolutional neural networks (GCNs), allowing the network to attend to different neighbors’ features with learnable weights, which helps GATs achieve state-of-the-art performance on node classification tasks in an inductive setting (Veličkovic et al., 2018). In our application of GATs to snRNA-seq data, we follow the exposition of Veličković et al. We first learned an approximate kNN graph (k=30) based on cell-to-cell pairwise Euclidean distances in 100-dimensional PCA space. The input to the GAT is a set of node features, . = {ℎ_1_, . . . , ℎ*_N_*}, where *N* is the number of nodes, and each vector *hi* corresponds to range-scaled, filtered and normalized gene counts for a particular node, ℎ*_i_* ∈ [0,1]^%^, where F is the number of features, i.e., genes (F=26,374). Each GAT layer produced a new set of node features, .′ = {ℎ_1_′, . . . , ℎ*_N_*′}, where ℎ*_i_*′ ∈ ℝ^F′^with cardinality *F*′.

To obtain .*h*′, first, a linear transformation was applied across nodes with a weight matrix, *W* ∈ ℝ*^F^*^′×^*^F^*. GAT layers differ from graph convolutional layers by including an attention mechanism. To calculate self-attention on nodes, we used a single-layer feedforward neural network with non-linear activations (LeakyReLU), yielding a single coefficient a: ℝ*^F^*^′^× ℝ*^F^*^′^→ ℝ per edge, *e_ij_* = *a*(*Wh_i_, Wh_j_*);ABℎ_!_, Bℎ_(_ C. These coefficients were normalized via a softmax function across all of neighbors of node *i*, 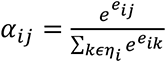 where η*_i_* tracks the neighborhood of node *i*; in our sparse implementation, we only considered the first-order neighborhood of each node. With these normalized attention coefficients, we can compute the linear combination of features to get the new node representation, 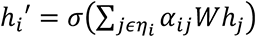 where σ is the logistic sigmoid function. We used multi-head attention and, prior to the final prediction layer, the new node feature representations were concatenated across heads and passed to the next GAT layer. In the final prediction layer, the node representations were averaged over the heads and a final non-linearity was applied (logistic sigmoid), 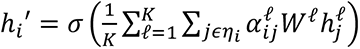 for K attention heads. Exponentiating the output of this final layer gave the predicted probability that a given node belongs to a particular class.

We used GATs to train 7 different models: 5 models that learn to discriminate SCA1 at each timepoint from other cells, 1 model to discriminate transcriptional signatures of SCA1 across mouse development (these 6 models comprise a binary, node classification task), and a final model to learn to identify each cell’s genotypic and developmental transcriptomic signature (1 model, 10 class labels). We randomly assigned cells to a training, validation, and test set to acquire three datasets such that *D_train_* ∩ *D_val_* ∩ *D_test_* = ∅, where each is a subset of the full mouse snRNA-seq dataset *D*, comprising 33%, 7%, and 7% of the total number of cells, respectively, formulating our tasks in an inductive setting (where the deep learning model has not seen the evaluation set graphs). We allowed each model to train for a maximum of 2,000 epochs, but set a patience of 100 epochs, evaluated on the validation set. The parameters were trained according to the objective function, which minimizes the negative log likelihood loss, with Adagrad optimization (Duchi et al., 2011; Glorot and Bengio, 2010). We used Glorot initialization for the weight matrices, *W*^ℓ^. For all models we used 2 layers with a hidden unit size of F’=8 across K=8 attention heads. We used cosine decay with 1000 first decay steps as our learning rate scheduler and a weight decay of 5e-3 for regularization. We used 50% dropout with alpha=0.2 as the slope in the LeakyReLU activation. We used Cluster-GCN to break up our kNN graph into sub-graphs with 5,000 partitions, facilitating mini-batching and stochastic gradient descent (Chiang et al., 2019). To evaluate our models, we report accuracy, sensitivity, and specificity at a threshold identified by Youden’s J-index for binary classification tasks (maximizing sensitivity and specificity). For multi-class problems, we report evaluation metrics with a one-vs.-rest heuristic evaluation scheme and take a prevalence-weighted averages such that each metric is adjusted for the number of class instances. Due to class imbalance and the desire to ensure our models learn interesting transcriptional signatures for distinguishing SCA1 at each time point (which comprise ∼10% of each dataset), we also report areas under the Receiving Operator Characteristic curve and Precision-Recall curve for each model. To implement and evaluate all models, we used PyTorch (version 1.6.0) and PyTorch Geometric (version 1.6.1) on a Nvidia GTX 1080-Ti GPU with CUDA v10.1 and 64-GB of CPU memory for graph pre-processing.

Given the excellent performance of our models across classification tasks, we sought to determine which transcriptional signatures our black-box models used to predict various labels, which allowed us to generate hypotheses as to the molecular mechanisms of SCA1 across brain development and aging. To identify the molecular determinants of SCA1 at a particular timepoint, given the appropriate model, we first looked at what transcripts the model highly weighted in making its decision, similar to saliency maps used in interpreting image classifiers (Simonyan et al., 2014). We extracted the learned weight matrix, 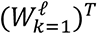, where k=1 indicates that the weight matrix is taken from the first layer for a given attention head and 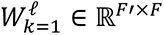 . In this case, the transpose of the learned matrix was a 26,374 x 8 matrix. To identify highly weighted genes from 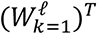, we sorted the absolute value of the weights per head and concatenated the result across heads, taking the union of weights up to an indexing variable for *m* top genes, 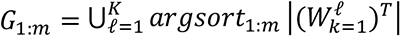. For each model, these weights can be found in and thought of as the transcripts important to discriminating between classes.

To visualize the important genes, we took the gene identities and visualized standardized expression across subsets of interest (cell type, genotype, and timepoint), representing expression with an average z-score from each subset (removing the mean from filtered and normalized expression values and dividing by the standard deviation across *D_test_* ). This amounted to a single-layer attribution method, which, although has been previously used with single-cell data, may omit contribution of downstream layers or interactions between layers in attributing the output of the deep network to input features (Sundararajan et al., 2018). In addition, looking at gene saliency globally may omit subtle differences between different types of cells. To simultaneously overcome both limitations, we also used the local interpretability approach of Integrated Gradients, which was developed around the axioms of sensitivity and implementation invariance (Sundararajan et al., 2018). Integrated Gradients assign an importance score to each input feature, i.e., each gene, by summing the gradients across the computation graph for an input, with respect to baseline, which we define to be no expression. For single-cell data and our GAT models, this amounted to 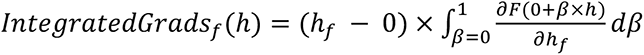, where F is a function that represents our deep network, 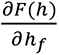 is the gradient of *F*(ℎ) along the gene feature dimension, *f*, and *h* is a vector in which filtered and normalized gene expression is averaged over cells of the same cell type, genotype, and timepoint and range-scaled, ℎ*_i_* ∈ [0,1]^F^. To smooth this gradient-based sensitivity map of output from features, we applied the SmoothGrad to our Integrated Gradients attribution approach (Smilkov et al., 2017). Using a SmoothGrad noise tunnel, we computed the attribution N=10 times, adding Gaussian noise, *N*(0,1), each iteration to each input *h*. In the SmoothGrad approach, the mean of the sampled attributions is returned as the importance score, which effectively smooths the attribution vector with a Gaussian Kernel. We finalized the importance score as the absolute value of the smoothed Integrated Gradients per feature and input, 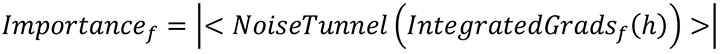, which yielded an importance score for each gene feature and averaged expression input, i.e., 13 cell types x 2 genotypes x 5 timepoints x 26,734 gene features or ∼3.4 million importance scores per model. To implement this attribution approach, we used PyTorch Captum (version 0.2.0) and custom functions to adapt these attribution methods to GNNs. Finally, to potentially identify unique, phenotypic populations, and to segregate single-cell embeddings by genotype and timepoint, we used PHATE and the attention coefficients from our 10-label GAT model to visualize the new graphs learned by each attention head. We constructed a matrix in which the *i*-th and *j*-th entry is given by 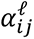, the normalized attention coefficients described above for a particular attention head.

These matrices are normalized adjacency matrices, yielding a directed graph. This graph can then be fed as a precursor to popular manifold learning methods such as UMAP and PHATE. The new low-dimensional embeddings learned by PHATE after inputting the graphs learned by various attention heads tend to segregate individual cells by the label, as each attention head learns a pattern in accordance with optimization of the objective function. To facilitate the use of state-of-the-art deep learning approaches for scientific interpretability and hypothesis-generation with other single-cell datasets, we have made all code publicly available for these deep learning modeling and interpretability approaches (see link in the Data Availability section).

### Statistical Analysis

Statistical analysis of snRNA-seq data was performed using Python (version 3.8.2), numpy (version 1.19.1), scipy (version 1.5.2), and various other open-source packages, described in further detail in the relevant sections. All other data were analyzed with Prism 8 (GraphPad) by two-tailed, unpaired Student’s t-test (comparing two experimental groups), one-way analysis of variance (ANOVA) (more than two experimental groups), or Kolmogorov-Smirnov tests (distributions of data) to assess statistical significance, unless otherwise noted. *P*<0.05 was considered to be statistically significant.

### Supplementary Figures

**Figure S1.**
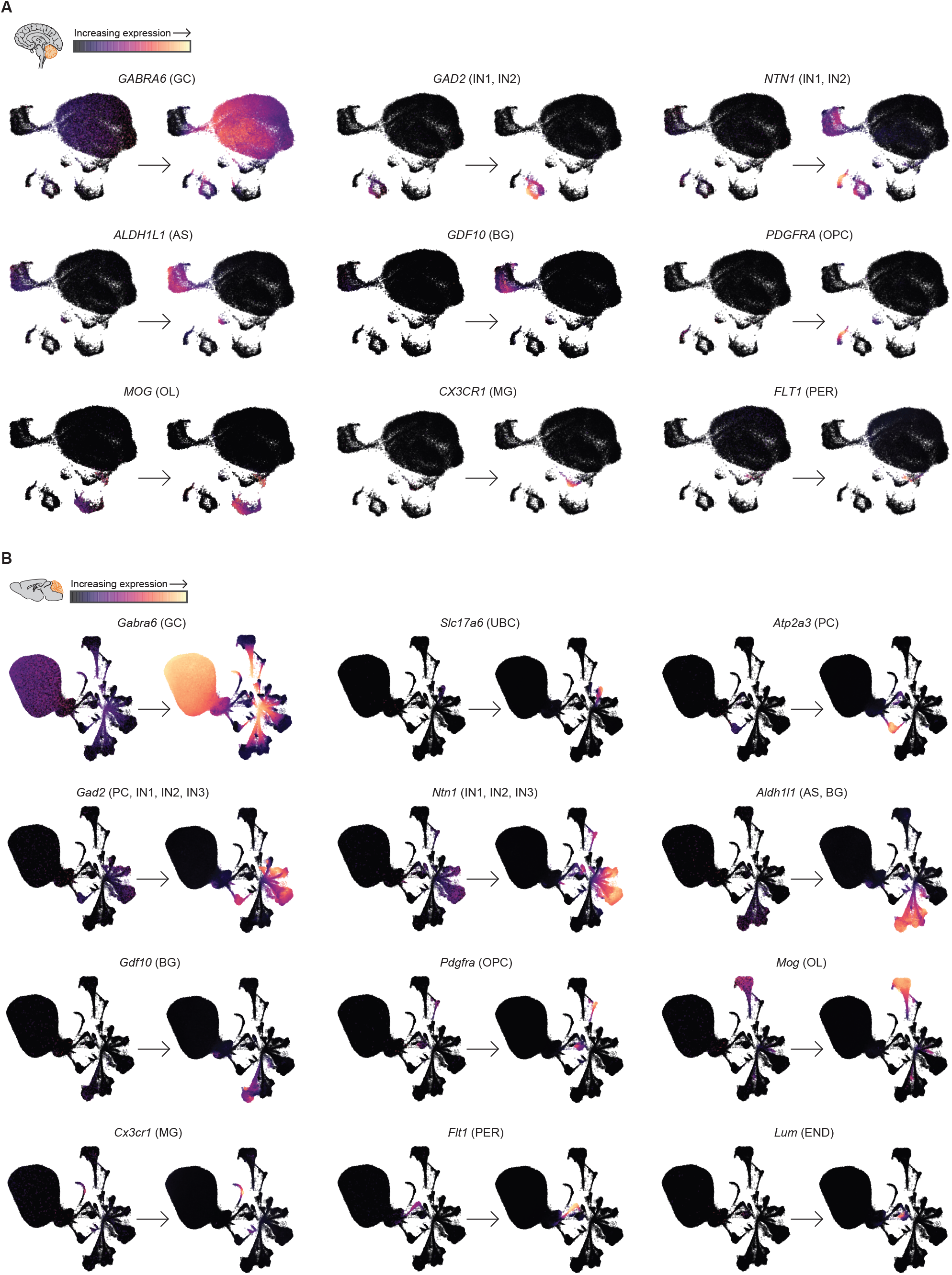
Data imputation using MAGIC improves data discriminability, *Related to Figure 1*. **(A and B)** UMAP embeddings of human (**A**) and mouse (**B**) snRNA-seq data before and after data imputation using Markov affinity-based graph imputation of cells (MAGIC), demonstrating improved select marker gene detection and cluster identification of imputed data (GC= granule cell, UBC= unipolar brush cell, PC= Purkinje cell, IN1= GABAergic interneuron 1, IN2= GABAergic interneuron 2, IN3= GABAergic interneuron 3, AS= astrocyte, BG= Bergmann glia, OPC= oligodendrocyte progenitor cell, OL= oligodendrocyte, MG= microglia, PER= pericyte, END= endothelial cell). Nuclei were extracted from the cerebellum of 6 human post-mortem samples (41,150 nuclei) and 30 mice from 5, 12, 18, 24, and 30 weeks of age (318,312 nuclei).

**Figure S2.**
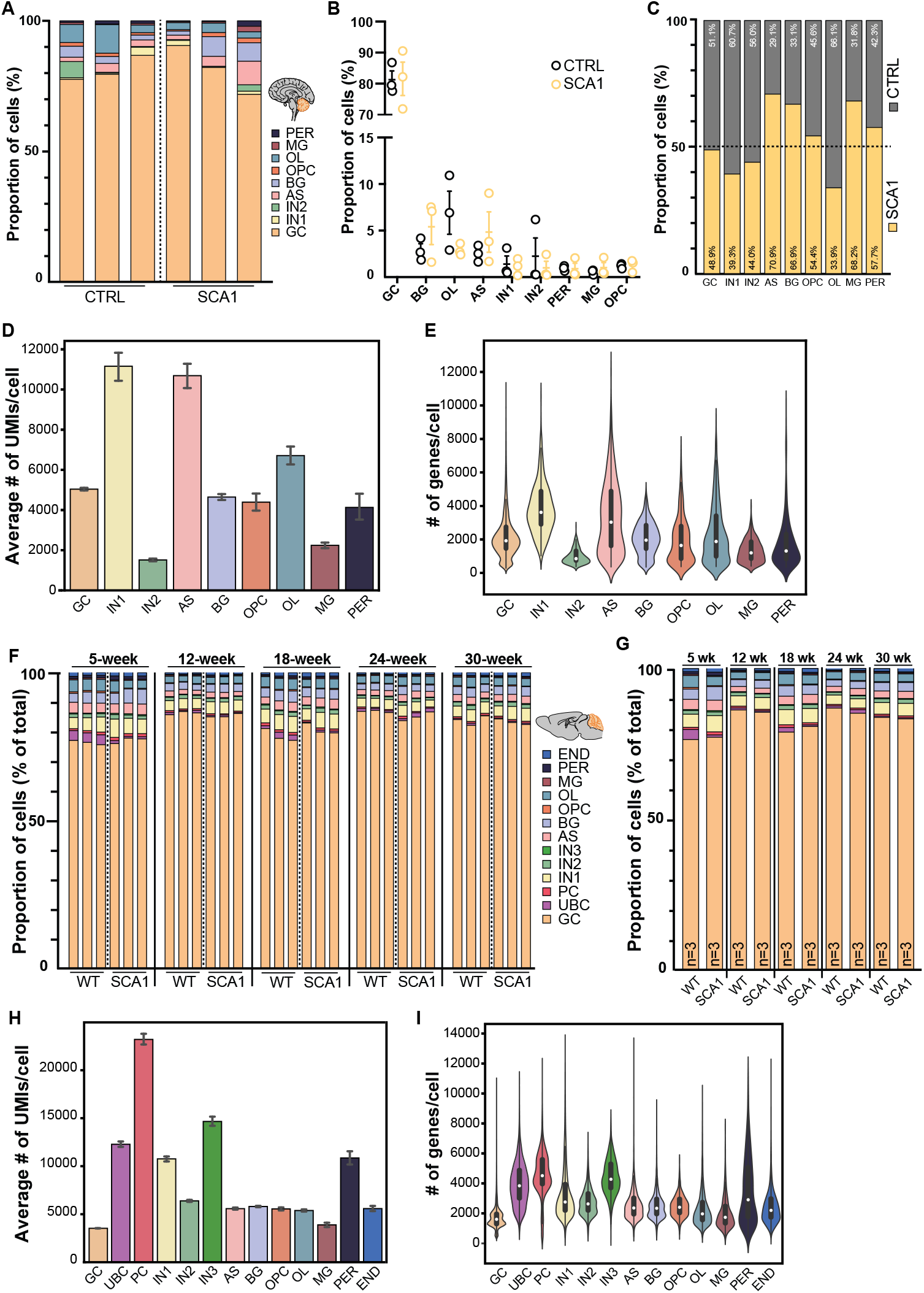
Representation of different cell types across samples and quality control information for clusters, *Related to Figure 1*. **(A)** Relative distribution of cell types in individual human samples. **(B)** Comparison of cluster proportions (% of total) between CTRL and SCA1. Data shown are mean±s.e.m. Student’s t-tests were used to compare across genotypes within a cluster and no significant differences were observed (CTRL, n=3; SCA1, n=3 patients). **(C)** Contribution of CTRL and SCA1 samples to each cluster after normalization to control for differences in total number of cells across genotypes (CTRL, n=3; SCA1, n=3 patients). **(D)** Bar plots of number of unique molecular identifiers (UMIs) detected per cell in each human cluster. Mean and 95% confidence interval are displayed. **(E)** Violin plots of number of genes detected per cell in each human cluster before imputation with MAGIC. Data shown are median (white dot) and interquartile range. **(F and G)** Relative distributions of cell types in the mouse dataset by individual sample (F) or grouped by condition after merging biological replicates (G). **(H)** Bar plots of number of UMIs detected per cell in each mouse cluster. Mean and 95% confidence interval are displayed. **(I)** Violin plots of number of genes detected per cell in each mouse cluster before imputation with MAGIC. Data shown are median (white dot) and interquartile range.

**Figure S3.**
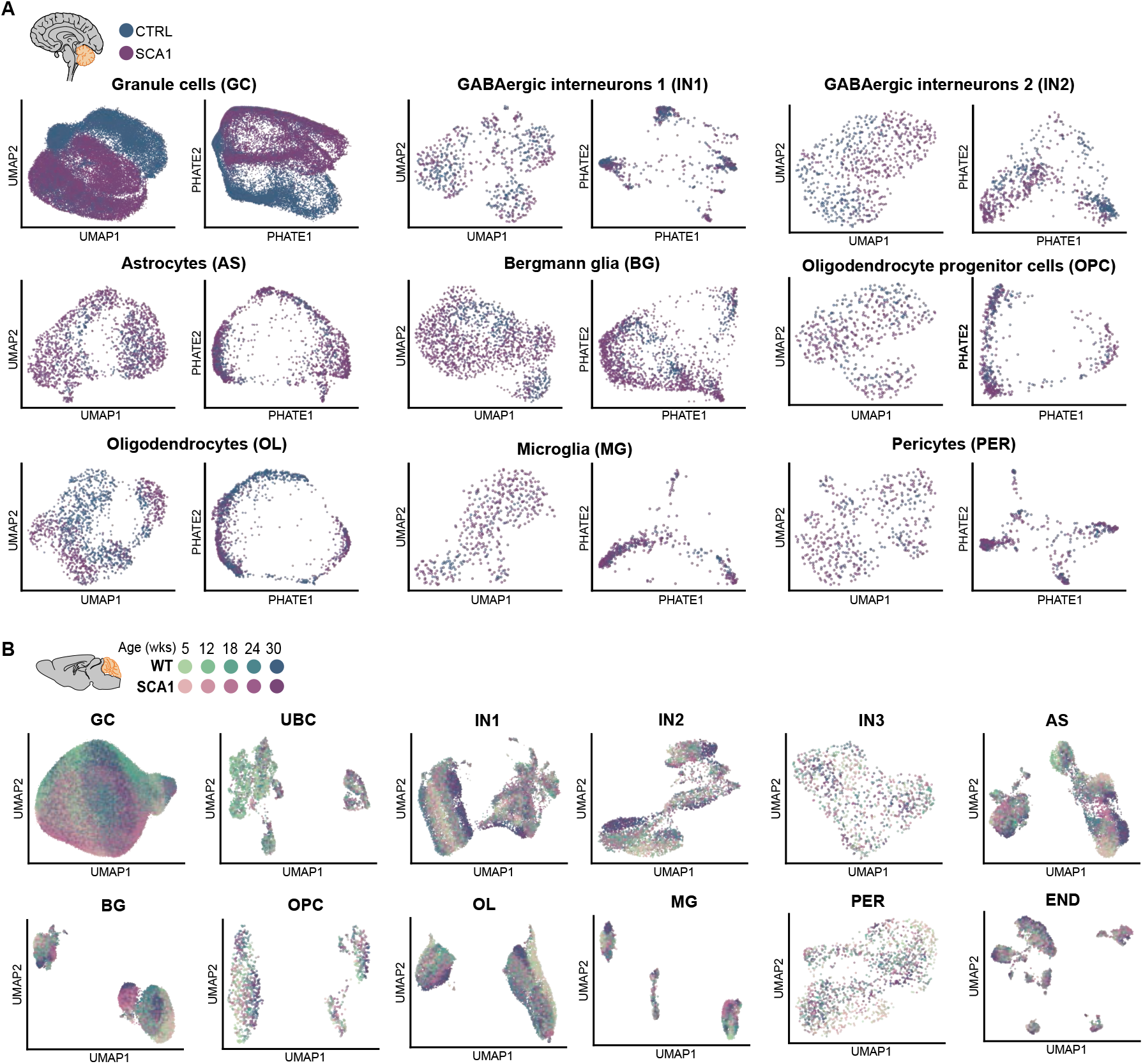
Visualization of individual cell types to identify sub-clusters, *Related to Figure 2*. **(A)** UMAP and PHATE embeddings calculated for individual clusters in the human datasets (GC= 14,043 CTRL and 19,313 SCA1 nuclei; IN1= 349 CTRL and 326 SCA1 nuclei; IN2= 312 CTRL and 353 SCA1 nuclei; AS= 408 CTRL and 1,430 SCA1 nuclei; BG= 437 CTRL and 1,272 SCA1 nuclei; OPC= 184 CTRL and 316 SCA1 nuclei; OL= 871 CTRL and 643 SCA1 nuclei; MG= 106 CTRL and 307 SCA1 nuclei; PER= 155 CTRL and 305 SCA1 nuclei). **(B)** UMAP embedding calculated for individual cell types in the mouse dataset to identify sub-clusters (GC= 134,797 WT and 126,067 SCA1 nuclei; UBC= 2,233 WT and 919 SCA1 nuclei; PC= 1,152 WT and 1,026 SCA1 nuclei; IN1= 6,797 WT and 6,710 SCA1 nuclei; IN2= 1,968 WT and 1,907 SCA1 nuclei; IN3= 457 WT and 433 SCA1 nuclei; AS= 4,088 WT and 4,097 SCA1 nuclei; BG= 4,825 WT and 4,771 SCA1 nuclei; OPC= 578 WT and 430 SCA1 nuclei; OL= 5,118 WT and 4,267 SCA1 nuclei; MG= 578 WT and 546 SCA1 nuclei; PER= 756 WT and 752 SCA1 nuclei; END= 1,375 WT and 1,665 SCA1 nuclei). Data are colored by timepoint (mouse) and genotype (mouse and human).

**Figure S4.**
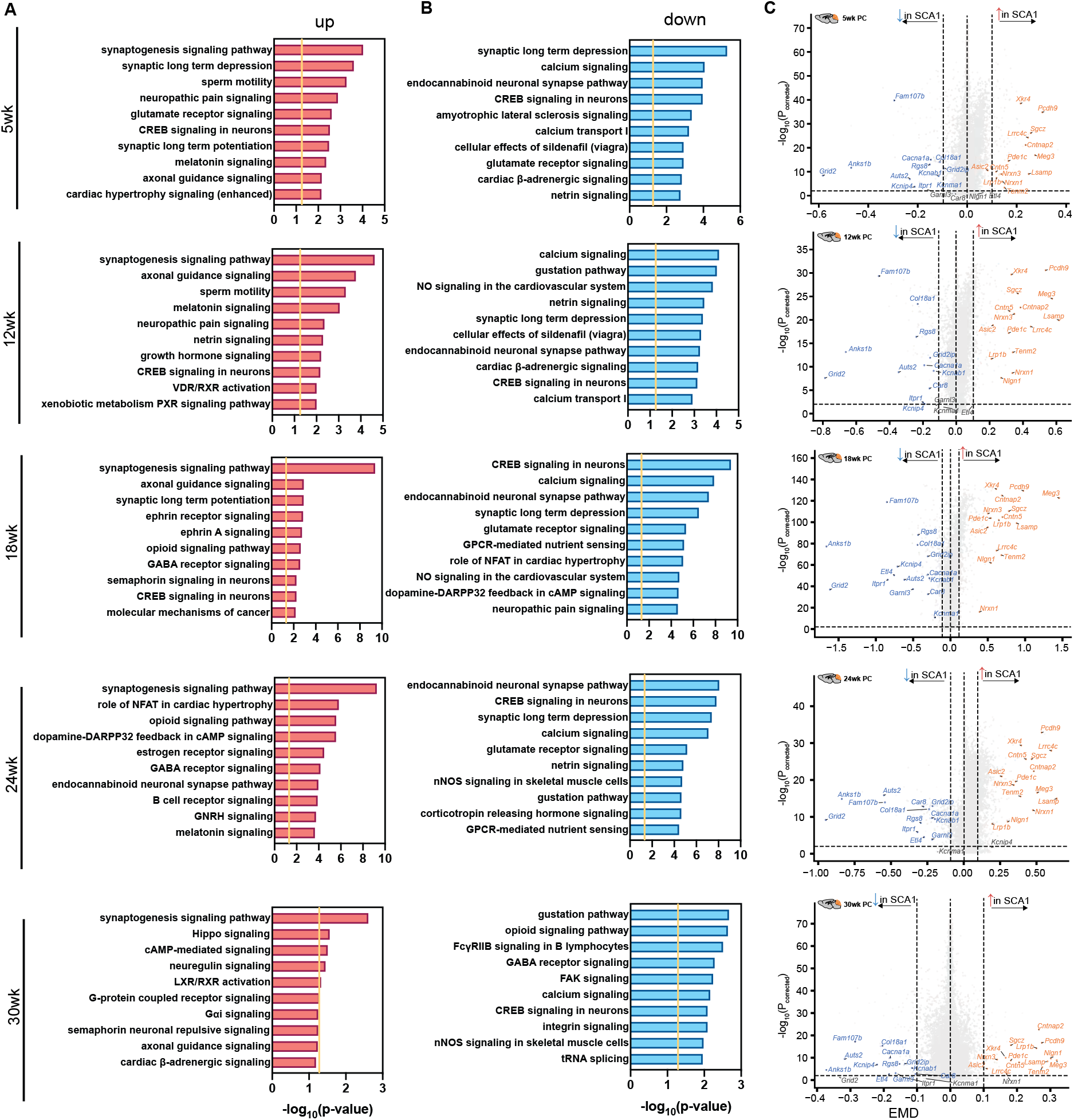
Purkinje cell dynamics over time, *Related to Figure 2*. **(A and B)** Top 10 enriched ingenuity canonical pathways from Ingenuity Pathway Analyses of up- (A) and down-regulated (B) DEGs in PCs at 5, 12, 18, 24, 30 weeks of age. Genes with imputed |EMD|≥0.1 and P_corrected_<0.01 were considered significant. Significant pathways were determined by a cut-off of -log(P-value)>1.3. **(C)** Volcano plot displaying several significant DEGs of interest of the mouse PC cluster by timepoint (5-week, 92 up- and 82 down-regulated genes; 12-week, 219 up- and 115 down- regulated genes; 18-week, 784 up- and 278 down-regulated genes; 24-week, 1,052 up- and 177 down-regulated genes; 30-week, 63 up- and 101 down-regulated genes).

**Figure S5.**
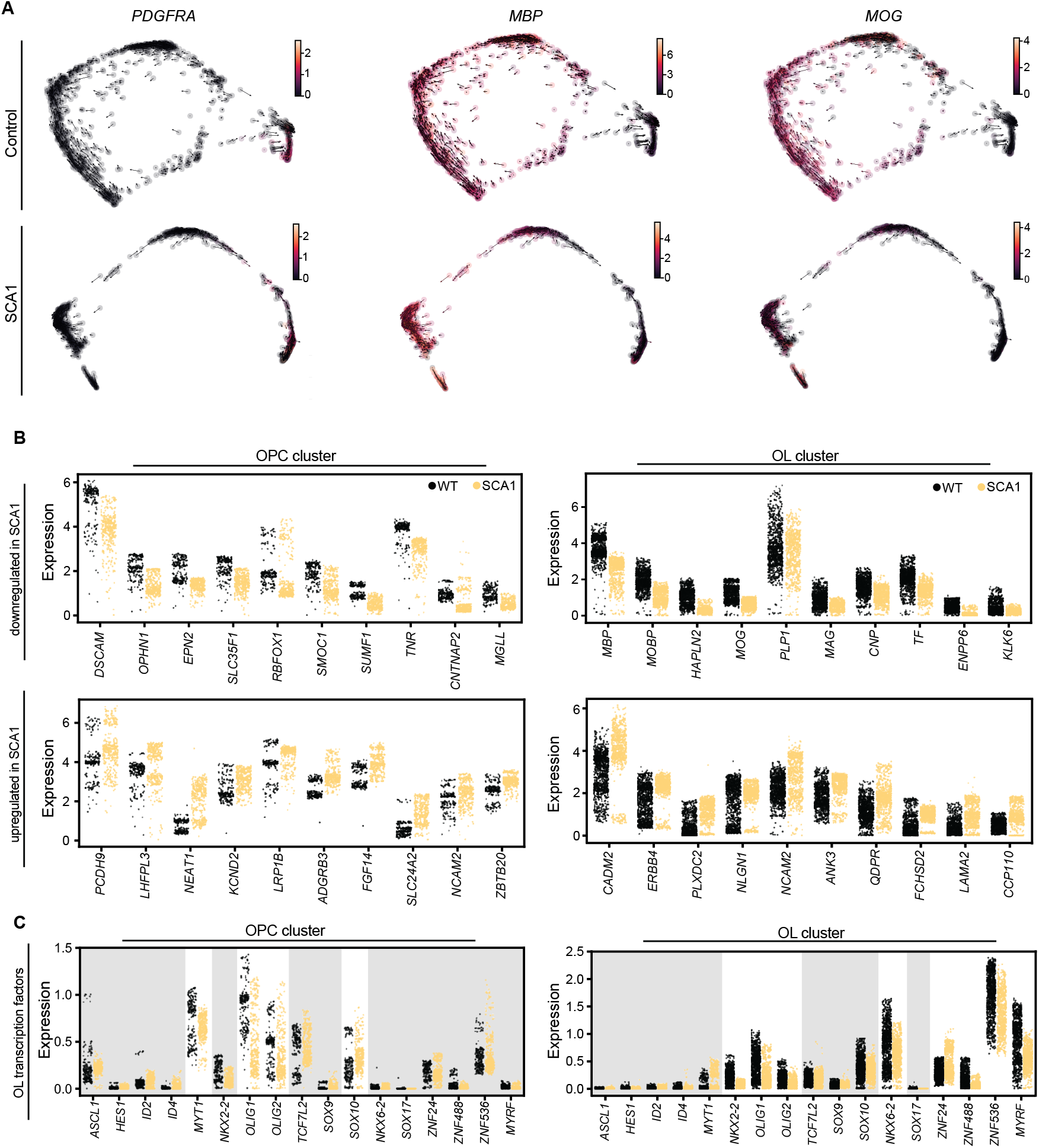
Analysis of the human oligodendroglia lineage with snRNA-seq, *Related to Figure 3*. **(A)** RNA velocity of OPC marker *PDGFRA* and OL markers *MBP* and *MOG* inferred by scVelo and projected onto PHATE embeddings of human oligodendroglia (OPC+OL clusters; CTRL, n=1,055; SCA1, n=959 nuclei). **(B)** Jitter plots of human OPC (left) and OL (right) clusters displaying distribution of expression of selected down- (top) and up-regulated (bottom) genes. **(C)** Jitter plots of human OPC (left) and OL (right) clusters displaying distribution of expression of OL-related transcription factors. Genes with imputed |EMD|≥0.1 and P_corrected_<0.01 were considered significant. Genes with gray shading represent non-significant changes.

**Figure S6.**
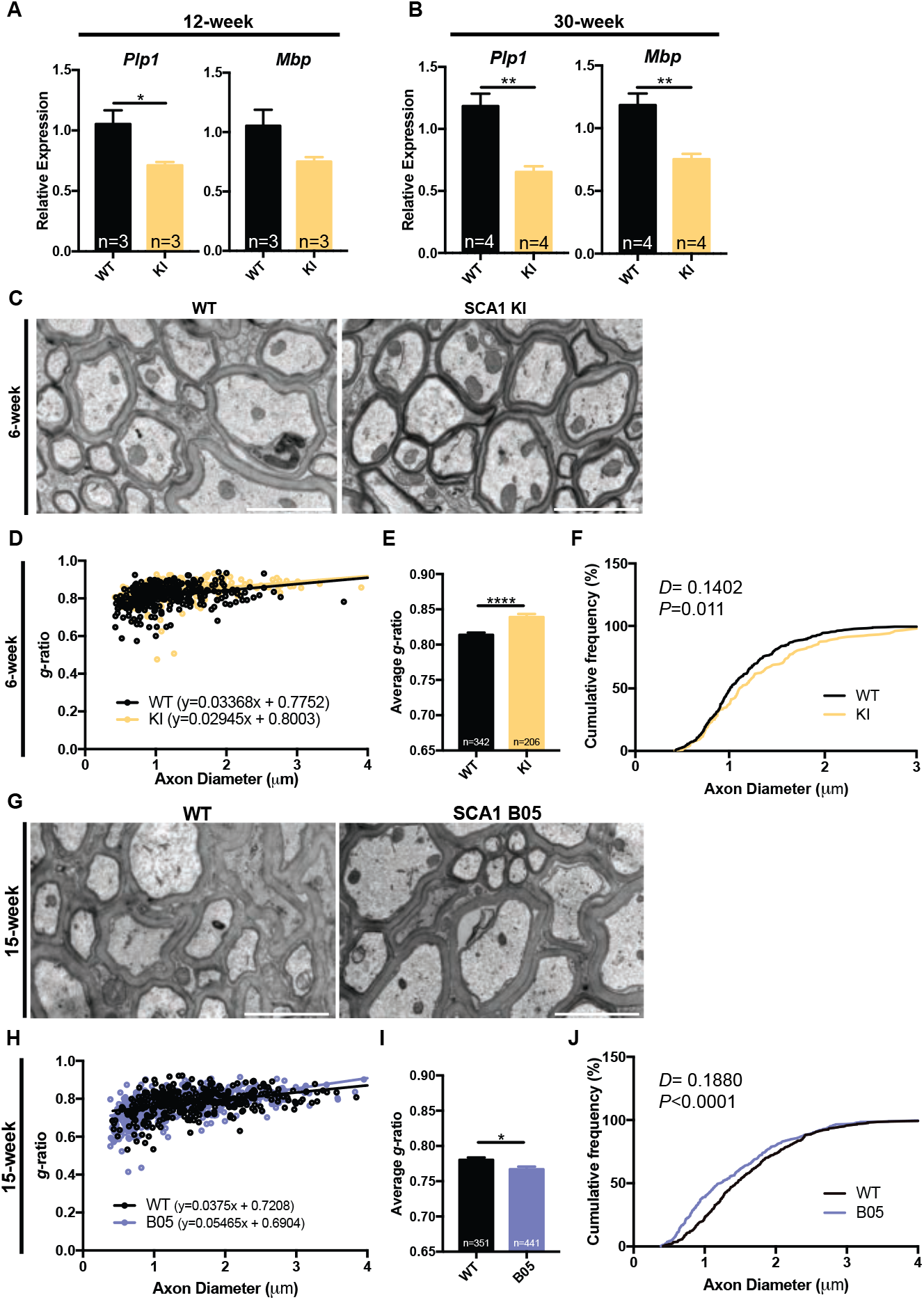
Early myelination deficits in SCA1 are progressive and not secondary to Purkinje cell dysfunction, *Related to Figure 3*. **(A and B)** qRT-PCR validation of bulk RNA-seq changes for myelin transcripts *Mbp* and *Plp1* in the 12-week (A) and 30-week (B) SCA1 KI mouse cerebellum (12-week: WT, n=3; SCA1 KI, n=3 mice; 30-week: WT, n=4; SCA1 KI, n=4 mice). **(C)** Representative transmission electron microscopy images of myelinated PC axons from 6-week WT and SCA1 KI mouse cerebellum. **(D-F)** Quantification of *g*-ratio (D,E) and axon diameter (F) from 6-week WT and SCA KI mice in (C), showing a slight decrease in myelination in SCA KI mice (WT, n=1; SCA1 KI, n=1 mouse). Scale bar=2μm. **(G)** Representative transmission electron microscopy images of myelinated Purkinje cell axons from 15-week WT and SCA1 PC-specific transgenic B05 (*Pcp2-ATXN1[82Q]/+*) mouse cerebellum. **(H-J)** Quantification of *g*-ratio (H,I) and axon diameter (J) from 15-week WT and SCA1 B05 mice in (G), showing a subtle increase in myelination (WT, n=2; SCA1 B05, n=2 mice). Scale bar=2μm. Only myelinated axon diameters were included in (F and J). For bar plots, data presented are mean±s.e.m. The following statistical tests were used: Student’s t-tests to compare across genotypes for (A, B, E, and I); Kolmogorov-Smirnov test to compare frequency distributions in (F and J). \**P*<0.05, \*\**P*<0.01, \*\*\*\**P*<0.0001.

**Figure S7.**
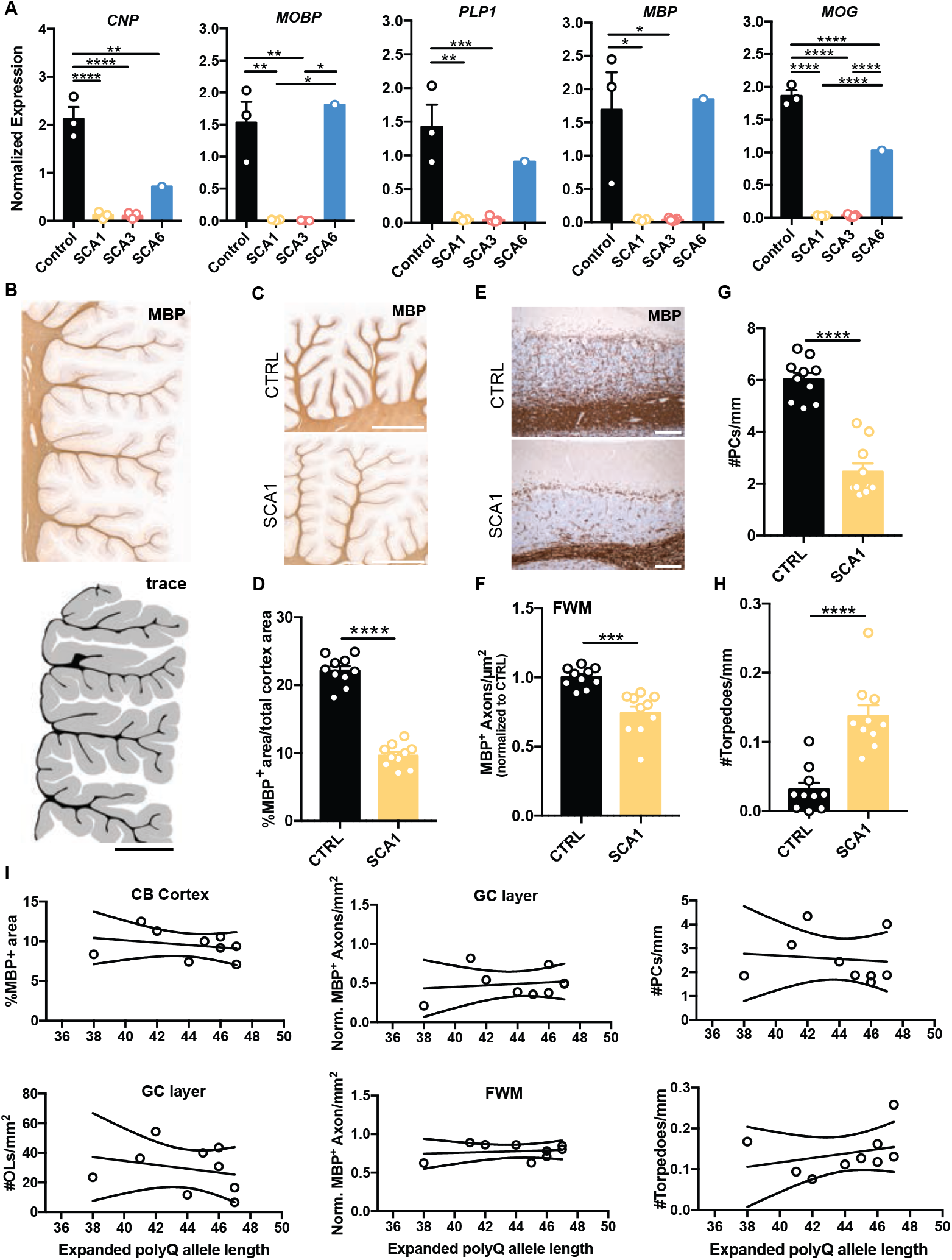
Validation of myelin deficits in human SCA post-mortem cerebellum, *Related to Figure 3*. **(A)** qRT-PCR validation of myelin gene down-regulation in SCA1 and SCA3, but not SCA6 human post-mortem cerebellar (CB) cortex (CTRL, n=3; SCA1, n=3; SCA3, n=5; SCA6, n=1 patient). **(B)** Representative images MBP immunostaining of the human post-mortem cerebellum and tracing to determine % of MBP-positive area in the CB cortex. Scale bar=4mm. **(C and D)** Representative images (C) and quantification (D) of % of MBP-positive area in the total cerebellar cortex (CTRL, n=10; SCA1, n=10 patients). Scale bar=4mm. **(E and F)** Representative images (E) and quantification (F) of MBP-positive axon profiles in the folial white matter (FWM) (CTRL, n=10; SCA1, n=10 patients). Scale bar=100μm. **(G and H)** Quantification of PCs (G) and PC axon torpedoes (H) in human post-mortem tissue (CTRL, n=10; SCA1, n=10 patients). **(I)** Relationship between *ATXN1* polyQ length and measured pathological changes in the granule cell layer (GC layer), the CB cortex, and folial WM, showing a trending decrease in overall myelin area and number of OLs with higher repeat lengths in SCA1. For bar plots, data are mean±s.e.m. For line plots of polyQ length-dependence, 95% confidence intervals are displayed. The following statistical tests were used: one-way ANOVA with multiple comparisons to compare across genotypes for (A); Student’s t-tests to compare across genotypes for (D, F G, and H). \**P*<0.05, \*\**P*<0.01, \*\*\**P*<0.001, \*\*\*\**P*<0.0001, n.s.=not significant.

**Figure S8.**
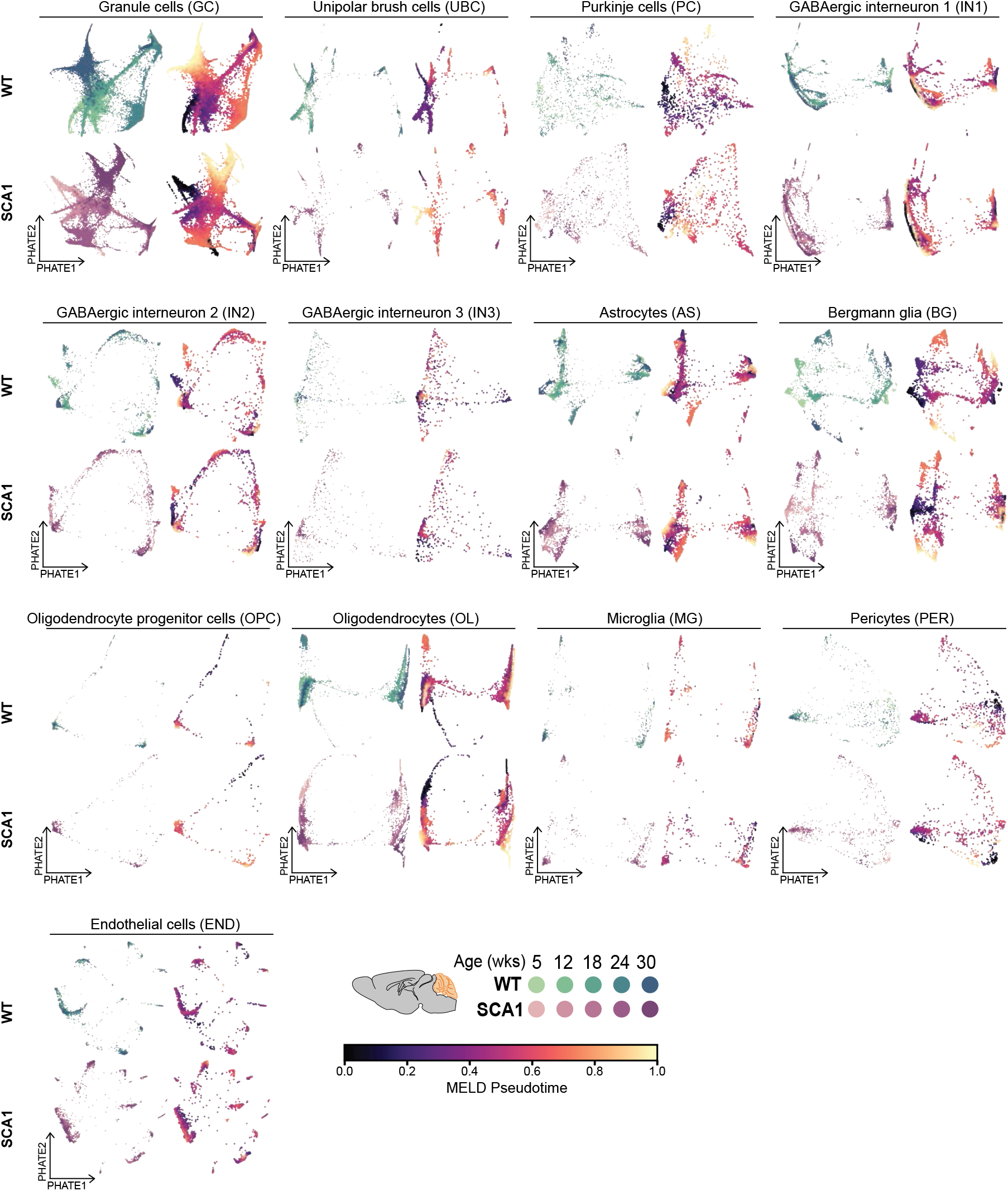
Comparison of biological time series data with pseudotemporally-ordered continuous cell trajectories, *Related to Figure 4*. PHATE embeddings calculated for individual cell types in the mouse dataset colored by biological time (left) and MELD pseudotime (right) for WT (top) and SCA1 (bottom) cells to confirm accurate pseudotemporal ordering of cells.

### Supplementary Tables

**Table S1. Details of human post-mortem cerebellum specimens, Related to Figure 1**

Details of control (CTRL) and SCA patient cerebellar cortex samples used for each experiment (PMI=post-mortem interval, snRNA-seq=single-nucleus RNA-seq, qRT-PCR=quantitative RT-PCR, WB=western blotting, IHC=immunohistochemistry, n.d.=not determined).

**Table S2. Tissue information and number of cells sequenced for mouse and human snRNA-samples, *Related to Figure 1***

**(A and B)** Information regarding sample ID, age, genotype, sex, and number of cells for each mouse (A) and human cerebellar cortex (B) sample sequenced.

**Table S3. Number of differentially expressed genes in each cell type at each timepoint, Related to Figure 1**

**(A)** Number of up- and down-regulated genes in each of the 13 SCA1 KI mouse cerebellar cell types at 5, 12, 18, 24, and 30 weeks of age.

**(B)** Number of up- and down-regulated genes in each of the 9 human SCA1 cerebellar cell types in post-mortem tissue.

GC= granule cells, UBC= unipolar brush cells, PC= Purkinje cells, IN1= GABAergic interneuron 1, IN2= GABAergic interneuron 2, IN3= GABAergic interneuron 3, AS= astrocyte, BG= Bergmann glia, OPC= oligodendrocyte progenitor cells, OL= oligodendrocyte, MG= microglia, PER= pericyte, END= endothelial cell.

**Table S4. Imputed differential gene expression between CTRL and SCA1 human post-mortem cerebellum, *Related to Figure 1***

Lists of up- and down-regulated genes in each of the 9 human cerebellar cell types (after imputation within genotype) organized by sheet. EMD was calculated to compare distributions and |EMD|≥0.1 and Benjamini-Hochberg adjusted P-value (P<0.01) were used to determine significance.

**Table S5. Imputed differential gene expression between WT and SCA1 KI mouse cerebellum, *Related to Figure 1***

Lists of up- and down-regulated genes in each of the 13 mouse cerebellar cell types (after imputation within genotype) organized by sheet. EMD was calculated to compare distributions and |EMD|≥0.1 and Benjamini-Hochberg adjusted P-value (P<0.01) were used to determine significance.

**Table S6. Gene ontology of differentially expressed genes in SCA1 KI mouse Purkinje cells, Related to Figure 2**

Lists of enriched ingenuity canonical pathways from lists of all DEGs (up and down), up-regulated genes only, or down-regulated genes only at each timepoint in mouse PCs. Significant pathways were determined by a cut-off of -log(P-value)>1.3.

**Table S7. Expression of top dynamical genes for each cell type in the WT and SCA1 KI mouse cerebellum, *Related to Figure 4***

Lists of the top 100 dynamical genes for each genotype within each cell type. Expression differences for each gene were calculated between genotypes (SCA1-WT) for each pseudotime bin. Importance ratings refer to the information gain (improvement in accuracy to the branches that a gene feature is on).

**Table S8. Top salient genes for GAT, Related to Figure 4**

List of top 20 salient genes per GAT model for predicting disease status label at each timepoint based on ranking the first GAT layer’s learned weight matrix, transforming input features to their hidden unit representations.

**Table S9. Top important genes based on local interpretability from Integrated Gradients analysis, *Related to Figure 4***

List of top 50 most important genes the Integrated Gradients analysis with a SmoothGrad noise tunnel for each representative cell type and timepoint, based on average expression in each subset.

## Notes

### Competing Interest Statement

The authors have declared no competing interest.

## REFERENCES

Al-Dalahmah, O., Sosunov, A.A., Shaik, A., Ofori, K., Liu, Y., Vonsattel, J.P., Adorjan, I., Menon, V., and Goldman, J.E. (2020). Single-nucleus RNA-seq identifies Huntington disease astrocyte states. Acta Neuropathol Commun 8, 19.

Barnes, J.A., Ebner, B.A., Duvick, L.A., Gao, W., Chen, G., Orr, H.T., and Ebner, T.J. (2011). Abnormalities in the climbing fiber-Purkinje cell circuitry contribute to neuronal dysfunction in ATXN1[82Q] mice. J Neurosci 31, 12778–12789.

Barron, T., Saifetiarova, J., Bhat, M.A., and Kim, J.H. (2018). Myelination of Purkinje axons is critical for resilient synaptic transmission in the deep cerebellar nucleus. Sci Rep 8, 1022.

Becht, E., McInnes, L., Healy, J., Dutertre, C.-A., Kwok, I.W.H., Ng, L.G., Ginhoux, F., and Newell, E.W. (2018). Dimensionality reduction for visualizing single-cell data using UMAP. Nature Biotechnology 37, 38–44.

Bergen, V., Lange, M., Peidli, S., Wolf, F.A., and Theis, F.J. (2020). Generalizing RNA velocity to transient cell states through dynamical modeling. Nat Biotechnol.

Bergstra, J., Yamins, D., and Cox, D.D. (2013). Making a Science of Model Search: Hyperparameter Optimization in Hundreds of Dimensions for Vision Architectures. International Conference on Machine Learning, 9.

Burkhardt, D.B., Stanley, J.S., Tong, A., Perdigoto, A.L., Gigante, S.A., Herold, K.C., Wolf, G., Giraldez, A.J., van Dijk, D., and Krishnaswamy, S. (2020). Quantifying the effect of experimental perturbations in single-cell RNA-sequencing data using graph signal processing. bioRxiv, 532846.

Chen, T., and Guestrin, C. (2016). XGBoost: A Scalable Tree Boosting System (Association for Computing Machinery).

Chiang, W.-L., Liu, X., Si, S., Li, Y., Bengio, S., and Hsieh, C.-J. (2019). Cluster-GCN: An Efficient Algorithm for Training Deep and Large Graph Convolutional Networks (Association for Computing Machinery).

Choe, M., Cortes, E., Vonsattel, J.P., Kuo, S.H., Faust, P.L., and Louis, E.D. (2016). Purkinje cell loss in essential tremor: Random sampling quantification and nearest neighbor analysis. Mov Disord 31, 393–401.

Chopra, R., Bushart, D.D., Cooper, J.P., Yellajoshyula, D., Morrison, L.M., Huang, H., Scoles, D.R., Pulst, S.M., Orr, H.T., and Shakkottai, V.G. (2020). Altered Capicua expression drives regional Purkinje neuron vulnerability through ion channel gene dysregulation in Spinocerebellar ataxia type 1. bioRxiv, 2020.2005.2021.104976.

Cordell, H.J. (2009). Detecting gene-gene interactions that underlie human diseases. Nat Rev Genet 10, 392–404.

Costa, M.D.C., Radzwion, M., McLoughlin, H.S., Ashraf, N.S., Fischer, S., Shakkottai, V.G., Maciel, P., Paulson, H.L., and Oz, G. (2020). In Vivo Molecular Signatures of Cerebellar Pathology in Spinocerebellar Ataxia Type 3. Mov Disord.

Cvetanovic, M., Ingram, M., Orr, H., and Opal, P. (2015). Early activation of microglia and astrocytes in mouse models of spinocerebellar ataxia type 1. Neuroscience 289, 289–299.

Driessen, T.M., Lee, P.J., and Lim, J. (2018). Molecular pathway analysis towards understanding tissue vulnerability in spinocerebellar ataxia type 1. eLife 7.

Duchi, J., Hazan, E., and Singer, Y. (2011). Adaptive Subgradient Methods for Online Learning and Stochastic Optimization. Journal of Machine Learning Research 12, 2121–2159.

Duvick, L., Barnes, J., Ebner, B., Agrawal, S., Andresen, M., Lim, J., Giesler, G.J., Zoghbi, H.Y., and Orr, H.T. (2010). SCA1-like disease in mice expressing wild-type ataxin-1 with a serine to aspartic acid replacement at residue 776. Neuron 67, 929–935.

Ebner, B.A., Ingram, M.A., Barnes, J.A., Duvick, L.A., Frisch, J.L., Clark, H.B., Zoghbi, H.Y., Ebner, T.J., and Orr, H.T. (2013). Purkinje cell ataxin-1 modulates climbing fiber synaptic input in developing and adult mouse cerebellum. J Neurosci 33, 5806–5820.

Edamakanti, C.R., Do, J., Didonna, A., Martina, M., and Opal, P. (2018). Mutant ataxin1 disrupts cerebellar development in spinocerebellar ataxia type 1. Journal of Clinical Investigation 128, 2252–2265.

Fields, R.D. (2015). A new mechanism of nervous system plasticity: activity-dependent myelination. Nat Rev Neurosci 16, 756–767.

Friedrich, J., Kordasiewicz, H.B., O’Callaghan, B., Handler, H.P., Wagener, C., Duvick, L., Swayze, E.E., Rainwater, O., Hofstra, B., Benneyworth, M., et al. (2018). Antisense oligonucleotide-mediated ataxin-1 reduction prolongs survival in SCA1 mice and reveals disease-associated transcriptome profiles. JCI Insight 3.

Glorot, X., and Bengio, Y. (2010). Understanding the difficulty of training deep feedforward neural networks (JMLR Workshop and Conference Proceedings).

Grubman, A., Chew, G., Ouyang, J.F., Sun, G., Choo, X.Y., McLean, C., Simmons, R.K., Buckberry, S., Vargas-Landin, D.B., Poppe, D., et al. (2019). A single-cell atlas of entorhinal cortex from individuals with Alzheimer’s disease reveals cell-type-specific gene expression regulation. Nat Neurosci 22, 2087–2097.

Habib, N., Avraham-Davidi, I., Basu, A., Burks, T., Shekhar, K., Hofree, M., Choudhury, S.R., Aguet, F., Gelfand, E., Ardlie, K.*, et al.* (2017). Massively parallel single-nucleus RNA- seq with DroNc-seq. Nat Methods 14, 955–958.

Haghverdi, L., Lun, A.T.L., Morgan, M.D., and Marioni, J.C. (2018). Batch effects in single-cell RNA-sequencing data are corrected by matching mutual nearest neighbors. Nat Biotechnol 36, 421–427.

Hills, L.B., Masri, A., Konno, K., Kakegawa, W., Lam, A.T., Lim-Melia, E., Chandy, N., Hill, R.S., Partlow, J.N., Al-Saffar, M., et al. (2013). Deletions in GRID2 lead to a recessive syndrome of cerebellar ataxia and tonic upgaze in humans. Neurology 81, 1378–1386.

Huang, B., Wei, W., Wang, G., Gaertig, Marta A., Feng, Y., Wang, W., Li, X.-J., and Li, S. (2015). Mutant Huntingtin Downregulates Myelin Regulatory Factor-Mediated Myelin Gene Expression and Affects Mature Oligodendrocytes. Neuron 85, 1212–1226.

Ingram, M., Wozniak, E.A.L., Duvick, L., Yang, R., Bergmann, P., Carson, R., O’Callaghan, B., Zoghbi, H.Y., Henzler, C., and Orr, H.T. (2016). Cerebellar Transcriptome Profiles of ATXN1 Transgenic Mice Reveal SCA1 Disease Progression and Protection Pathways. Neuron 89, 1194–1207.

Jacobi, H., Reetz, K., du Montcel, S.T., Bauer, P., Mariotti, C., Nanetti, L., Rakowicz, M., Sulek, A., Durr, A., Charles, P., et al. (2013). Biological and clinical characteristics of individuals at risk for spinocerebellar ataxia types 1, 2, 3, and 6 in the longitudinal RISCA study: analysis of baseline data. The Lancet Neurology 12, 650–658.

Jafar-Nejad, P., Ward, C.S., Richman, R., Orr, H.T., and Zoghbi, H.Y. (2011). Regional rescue of spinocerebellar ataxia type 1 phenotypes by 14-3-3epsilon haploinsufficiency in mice underscores complex pathogenicity in neurodegeneration. Proc Natl Acad Sci U S A 108, 2142–2147.

Jakel, S., Agirre, E., Mendanha Falcao, A., van Bruggen, D., Lee, K.W., Knuesel, I., Malhotra, D., Ffrench-Constant, C., Williams, A., and Castelo-Branco, G. (2019). Altered human oligodendrocyte heterogeneity in multiple sclerosis. Nature 566, 543–547.

Kang, S.H., Li, Y., Fukaya, M., Lorenzini, I., Cleveland, D.W., Ostrow, L.W., Rothstein, J.D., and Bergles, D.E. (2013). Degeneration and impaired regeneration of gray matter oligodendrocytes in amyotrophic lateral sclerosis. Nat Neurosci 16, 571–579.

Kano, M., and Watanabe, T. (2017). Type-1 metabotropic glutamate receptor signaling in cerebellar Purkinje cells in health and disease. F1000Res 6, 416.

Kanton, S., Boyle, M.J., He, Z., Santel, M., Weigert, A., Sanchis-Calleja, F., Guijarro, P., Sidow, L., Fleck, J.S., Han, D., et al. (2019). Organoid single-cell genomic atlas uncovers human-specific features of brain development. Nature 574, 418–422.

Karadottir, R., Hamilton, N.B., Bakiri, Y., and Attwell, D. (2008). Spiking and nonspiking classes of oligodendrocyte precursor glia in CNS white matter. Nat Neurosci 11, 450–456.

Kashiwabuchi, N., Ikeda, K., Araki, K., Hirano, T., Shibuki, K., Takayama, C., Inoue, Y., Kutsuwada, T., Yagi, T., Kang, Y., et al. (1995). Impairment of motor coordination, Purkinje cell synapse formation, and cerebellar long-term depression in GluRo2 mutant mice. Cell 81, 245–252.

Kato, A.S., Knierman, M.D., Siuda, E.R., Isaac, J.T., Nisenbaum, E.S., and Bredt, D.S. (2012). Glutamate receptor delta2 associates with metabotropic glutamate receptor 1 (mGluR1), protein kinase Cgamma, and canonical transient receptor potential 3 and regulates mGluR1-mediated synaptic transmission in cerebellar Purkinje neurons. J Neurosci 32, 15296–15308.

Kim, J.H., Lukowicz, A., Qu, W., Johnson, A., and Cvetanovic, M. (2018). Astroglia contribute to the pathogenesis of spinocerebellar ataxia Type 1 (SCA1) in a biphasic, stage-of-disease specific manner. Glia 66, 1972–1987.

Koeppen, A.H. (2005). The pathogenesis of spinocerebellar ataxia. Cerebellum 4, 62–73.

Kohda, K., Kakegawa, W., Matsuda, S., Yamamoto, T., Hirano, H., and Yuzaki, M. (2013). The delta2 glutamate receptor gates long-term depression by coordinating interactions between two AMPA receptor phosphorylation sites. Proc Natl Acad Sci U S A 110, E948–957.

Korsunsky, I., Millard, N., Fan, J., Slowikowski, K., Zhang, F., Wei, K., Baglaenko, Y., Brenner, M., Loh, P.R., and Raychaudhuri, S. (2019). Fast, sensitive and accurate integration of single-cell data with Harmony. Nat Methods 16, 1289–1296.

La Manno, G., Soldatov, R., Zeisel, A., Braun, E., Hochgerner, H., Petukhov, V., Lidschreiber, K., Kastriti, M.E., Lonnerberg, P., Furlan, A., et al. (2018). RNA velocity of single cells. Nature 560, 494–498.

Lalouette, A., Guenet, J.L., and Vriz, S. (1998). Hotfoot mouse mutations affect the delta 2 glutamate receptor gene and are allelic to lurcher. Genomics 50, 9–13.

Lam, Y.C., Bowman, A.B., Jafar-Nejad, P., Lim, J., Richman, R., Fryer, J.D., Hyun, E.D., Duvick, L.A., Orr, H.T., Botas, J., et al. (2006). ATAXIN-1 interacts with the repressor Capicua in its native complex to cause SCA1 neuropathology. Cell 127, 1335–1347.

Lee, H., Fenster, R.J., Pineda, S.S., Gibbs, W.S., Mohammadi, S., Davila-Velderrain, J., Garcia, F.J., Therrien, M., Novis, H.S., Gao, F.*, et al.* (2020). Cell Type-Specific Transcriptomics Reveals that Mutant Huntingtin Leads to Mitochondrial RNA Release and Neuronal Innate Immune Activation. Neuron.

Lee, Y., Morrison, B.M., Li, Y., Lengacher, S., Farah, M.H., Hoffman, P.N., Liu, Y., Tsingalia, A., Jin, L., Zhang, P.W., et al. (2012). Oligodendroglia metabolically support axons and contribute to neurodegeneration. Nature 487, 443–448.

Lim, J., Crespo-Barreto, J., Jafar-Nejad, P., Bowman, A.B., Richman, R., Hill, D.E., Orr, H.T., and Zoghbi, H.Y. (2008). Opposing effects of polyglutamine expansion on native protein complexes contribute to SCA1. Nature 452, 713–718.

Lim, J., Hao, T., Shaw, C., Patel, A.J., Szabo, G., Rual, J.F., Fisk, C.J., Li, N., Smolyar, A., Hill, D.E., et al. (2006). A protein-protein interaction network for human inherited ataxias and disorders of Purkinje cell degeneration. Cell 125, 801–814.

Lin, S.C., Huck, J.H., Roberts, J.D., Macklin, W.B., Somogyi, P., and Bergles, D.E. (2005). Climbing fiber innervation of NG2-expressing glia in the mammalian cerebellum. Neuron 46, 773–785.

Lin, X., Antalffy, B., Kang, D., Orr, H.T., and Zoghbi, H.Y. (2000). Polyglutamine expansion down-regulates specific neuronal genes before pathologic changes in SCA1. Nat Neurosci 3, 157–163.

Luecken, M.D., Büttner, M., Chaichoompu, K., Danese, A., Interlandi, M., Mueller, M.F., Strobl, D.C., Zappia, L., Dugas, M., Colomé-Tatché, M., et al. (2020). Benchmarking atlas-level data integration in single-cell genomics. bioRxiv.

Lukas, C., Schols, L., Bellenberg, B., Rub, U., Przuntek, H., Schmid, G., Koster, O., and Suchan, B. (2006). Dissociation of grey and white matter reduction in spinocerebellar ataxia type 3 and 6: a voxel-based morphometry study. Neurosci Lett 408, 230–235.

Macosko, E.Z., Basu, A., Satija, R., Nemesh, J., Shekhar, K., Goldman, M., Tirosh, I., Bialas, A.R., Kamitaki, N., Martersteck, E.M., et al. (2015). Highly Parallel Genome-wide Expression Profiling of Individual Cells Using Nanoliter Droplets. Cell 161, 1202–1214.

Mandelli, M.L., De Simone, T., Minati, L., Bruzzone, M.G., Mariotti, C., Fancellu, R., Savoiardo, M., and Grisoli, M. (2007). Diffusion Tensor Imaging of Spinocerebellar Ataxias Types 1 and 2. American Journal of Neuroradiology 28, 1996–2000.

Martins Junior, C.R., Martinez, A.R.M., Vasconcelos, I.F., de Rezende, T.J.R., Casseb, R.F., Pedroso, J.L., Barsottini, O.G.P., Lopes-Cendes, I., and Franca, M.C., Jr. (2018). Structural signature in SCA1: clinical correlates, determinants and natural history. J Neurol 265, 2949–2959.

Mathys, H., Davila-Velderrain, J., Peng, Z., Gao, F., Mohammadi, S., Young, J.Z., Menon, M., He, L., Abdurrob, F., Jiang, X., et al. (2019). Single-cell transcriptomic analysis of Alzheimer’s disease. Nature 570, 332–337.

McKenzie, I.A., Ohayon, D., Li, H., de Faria, J.P., Emery, B., Tohyama, K., and Richardson, W.D. (2014). Motor skill learning requires active central myelination. Science 346, 318–322.

Moon, K.R., van Dijk, D., Wang, Z., Gigante, S., Burkhardt, D.B., Chen, W.S., Yim, K., Elzen, A.V.D., Hirn, M.J., Coifman, R.R., et al. (2019). Visualizing structure and transitions in high-dimensional biological data. Nat Biotechnol 37, 1482–1492.

Mot, A.I., Depp, C., and Nave, K.A. (2018). An emerging role of dysfunctional axon-oligodendrocyte coupling in neurodegenerative diseases. Dialogues Clin Neurosci 20, 283–292.

Nabavi, S., Schmolze, D., Maitituoheti, M., Malladi, S., and Beck, A.H. (2016). EMDomics: a robust and powerful method for the identification of genes differentially expressed between heterogeneous classes. Bioinformatics 32, 533–541.

Orlova, D.Y., Zimmerman, N., Meehan, S., Meehan, C., Waters, J., Ghosn, E.E., Filatenkov, A., Kolyagin, G.A., Gernez, Y., Tsuda, S., et al. (2016). Earth Mover’s Distance (EMD): A True Metric for Comparing Biomarker Expression Levels in Cell Populations. PLoS One 11, e0151859.

Orr, H.T., Chung, M.Y., Banfi, S., Kwiatkowski, T.J., Jr., Servadio, A., Beaudet, A.L., McCall, A.E., Duvick, L.A., Ranum, L.P., and Zoghbi, H.Y. (1993). Expansion of an unstable trinucleotide CAG repeat in spinocerebellar ataxia type 1. Nat Genet 4, 221–226.

Pan, M.K., Li, Y.S., Wong, S.B., Ni, C.L., Wang, Y.M., Liu, W.C., Lu, L.Y., Lee, J.C., Cortes, E.P., Vonsattel, J.G.*, et al.* (2020). Cerebellar oscillations driven by synaptic pruning deficits of cerebellar climbing fibers contribute to tremor pathophysiology. Sci Transl Med 12.

Polanski, K., Young, M.D., Miao, Z., Meyer, K.B., Teichmann, S.A., Park, J.-E., and Berger, B. (2019). BBKNN: fast batch alignment of single cell transcriptomes. Bioinformatics.

Power, E.M., Morales, A., and Empson, R.M. (2016). Prolonged Type 1 Metabotropic Glutamate Receptor Dependent Synaptic Signaling Contributes to Spino-Cerebellar Ataxia Type 1. J Neurosci 36, 4910–4916.

Ramani, B., Panwar, B., Moore, L.R., Wang, B., Huang, R., Guan, Y., and Paulson, H.L. (2017). Comparison of spinocerebellar ataxia type 3 mouse models identifies early gain-of-function, cell-autonomous transcriptional changes in oligodendrocytes. Hum Mol Genet 26, 3362–3374.

Ravindra, N., Sehanobish, A., Pappalardo, J.L., Hafler, D.A., and Dijk, D.V. (2020). Disease state prediction from single-cell data using graph attention networks. In Proceedings of the ACM Conference on Health, Inference, and Learning (Toronto, Ontario, Canada: Association for Computing Machinery), pp. 121–130.

Ruegsegger, C., Stucki, D.M., Steiner, S., Angliker, N., Radecke, J., Keller, E., Zuber, B., Ruegg, M.A., and Saxena, S. (2016). Impaired mTORC1-Dependent Expression of Homer-3 Influences SCA1 Pathophysiology. Neuron 89, 129–146.

Satija, R., Farrell, J.A., Gennert, D., Schier, A.F., and Regev, A. (2015). Spatial reconstruction of single-cell gene expression data. Nat Biotechnol 33, 495–502.

Schiebinger, G., Shu, J., Tabaka, M., Cleary, B., Subramanian, V., Solomon, A., Gould, J., Liu, S., Lin, S., Berube, P., et al. (2019). Optimal-Transport Analysis of Single-Cell Gene Expression Identifies Developmental Trajectories in Reprogramming. Cell 176, 928–943 e922.

Schirmer, L., Velmeshev, D., Holmqvist, S., Kaufmann, M., Werneburg, S., Jung, D., Vistnes, S., Stockley, J.H., Young, A., Steindel, M.*, et al.* (2019). Neuronal vulnerability and multilineage diversity in multiple sclerosis. Nature 573, 75–82.

Sehanobish, A., Ravindra, N., and van Dijk, D. (2020). Gaining insight into SARS-CoV-2 infection and COVID-19 severity using self-supervised edge features and Graph Neural Networks.

Serra, H.G., Byam, C.E., Lande, J.D., Tousey, S.K., Zoghbi, H.Y., and Orr, H.T. (2004). Gene profiling links SCA1 pathophysiology to glutamate signaling in Purkinje cells of transgenic mice. Hum Mol Genet 13, 2535–2543.

Servadio, A., Koshy, B., Armstrong, D., Antalffy, B., Orr, H.T., and Zoghbi, H.Y. (1995). Expression analysis of the ataxin-1 protein in tissues from normal and spinocerebellar ataxia type 1 individuals. Nat Genet 10, 94–98.

Shuvaev, A.N., Hosoi, N., Sato, Y., Yanagihara, D., and Hirai, H. (2017). Progressive impairment of cerebellar mGluR signalling and its therapeutic potential for cerebellar ataxia in spinocerebellar ataxia type 1 model mice. J Physiol 595, 141–164.

Simons, M., and Nave, K.A. (2015). Oligodendrocytes: Myelination and Axonal Support. Cold Spring Harb Perspect Biol 8, a020479.

Simonyan, K., Vedaldi, A., and Zisserman, A. (2014). Deep Inside Convolutional Networks: Visualising Image Classification Models and Saliency Maps.

Smilkov, D., Thorat, N., Kim, B., Viégas, F., and Wattenberg, M. (2017). SmoothGrad: removing noise by adding noise. International Conference on Machine Learning.

Stark, C., Breitkreutz, B.J., Reguly, T., Boucher, L., Breitkreutz, A., and Tyers, M. (2006). BioGRID: a general repository for interaction datasets. Nucleic Acids Res 34, D535–539.

Stoyas, C.A., Bushart, D.D., Switonski, P.M., Ward, J.M., Alaghatta, A., Tang, M.B., Niu, C., Wadhwa, M., Huang, H., Savchenko, A.*, et al.* (2020). Nicotinamide Pathway-Dependent Sirt1 Activation Restores Calcium Homeostasis to Achieve Neuroprotection in Spinocerebellar Ataxia Type 7. Neuron 105, 630–644 e639.

Stuart, T., Butler, A., Hoffman, P., Hafemeister, C., Papalexi, E., Mauck, W.M., 3rd, Hao, Y., Stoeckius, M., Smibert, P., and Satija, R. (2019). Comprehensive Integration of Single-Cell Data. Cell 177, 1888–1902 e1821.

Sundararajan, M., Taly, A., and Yan, Q. (2018). Axiomatic Attribution for Deep Networks. International Conference on Learning Represeations.

Tejwani, L., and Lim, J. (2020). Pathogenic mechanisms underlying spinocerebellar ataxia type 1. Cell Mol Life Sci.

Utine, G.E., Haliloglu, G., Salanci, B., Cetinkaya, A., Kiper, P.O., Alanay, Y., Aktas, D., Boduroglu, K., and Alikasifoglu, M. (2013). A homozygous deletion in GRID2 causes a human phenotype with cerebellar ataxia and atrophy. J Child Neurol 28, 926–932.

van Dijk, D., Sharma, R., Nainys, J., Yim, K., Kathail, P., Carr, A.J., Burdziak, C., Moon, K.R., Chaffer, C.L., Pattabiraman, D., et al. (2018). Recovering Gene Interactions from Single-Cell Data Using Data Diffusion. Cell 174, 716–729.e727.

Velickovic, P., Cucurull, G., Casanova, A., Romero, A., Lio, P., and Bengio, Y. (2018). Graph Attention Networks. International Conference on Learning Representations, 12.

Vonsattel, J.P., Amaya Mdel, P., Cortes, E.P., Mancevska, K., and Keller, C.E. (2008). Twenty-first century brain banking: practical prerequisites and lessons from the past: the experience of New York Brain Bank, Taub Institute, Columbia University. Cell Tissue Bank 9, 247–258.

Wang, T., and Nabavi, S. (2018). SigEMD: A powerful method for differential gene expression analysis in single-cell RNA sequencing data. Methods 145, 25–32.

Wang, Y., and Blei, D.M. (2020). The Blessings of Multiple Causes. Journal of the American Statistical Association 114, 1574–1596.

Watase, K., Weeber, E.J., Xu, B., Antalffy, B., Yuva-Paylor, L., Hashimoto, K., Kano, M., Atkinson, R., Sun, Y., Armstrong, D.L.*, et al.* (2002). A Long CAG Repeat in the Mouse Sca1 Locus Replicates SCA1 Features and Reveals the Impact of Protein Solubility on Selective Neurodegeneration. Neuron 34, 905–919.

Welch, J.D., Kozareva, V., Ferreira, A., Vanderburg, C., Martin, C., and Macosko, E.Z. (2019). Single-Cell Multi-omic Integration Compares and Contrasts Features of Brain Cell Identity. Cell 177, 1873–1887 e1817.

Wolf, F.A., Angerer, P., and Theis, F.J. (2018). SCANPY: large-scale single-cell gene expression data analysis. Genome Biol 19, 15.

Young, K.M., Psachoulia, K., Tripathi, R.B., Dunn, S.J., Cossell, L., Attwell, D., Tohyama, K., and Richardson, W.D. (2013). Oligodendrocyte dynamics in the healthy adult CNS: evidence for myelin remodeling. Neuron 77, 873–885.

Zhou, Y., Song, W.M., Andhey, P.S., Swain, A., Levy, T., Miller, K.R., Poliani, P.L., Cominelli, M., Grover, S., Gilfillan, S., et al. (2020). Human and mouse single-nucleus transcriptomics reveal TREM2-dependent and TREM2-independent cellular responses in Alzheimer’s disease. Nat Med 26, 131–142.

Zoghbi, H.Y., and Orr, H.T. (2009). Pathogenic mechanisms of a polyglutamine-mediated neurodegenerative disease, spinocerebellar ataxia type 1. J Biol Chem 284, 7425–7429.

Zonouzi, M., Scafidi, J., Li, P., McEllin, B., Edwards, J., Dupree, J.L., Harvey, L., Sun, D., Hubner, C.A., Cull-Candy, S.G., et al. (2015). GABAergic regulation of cerebellar NG2 cell development is altered in perinatal white matter injury. Nat Neurosci 18, 674–682.

